# Structural and functional characterization of SARS-CoV-2 RBD domains produced in mammalian cells

**DOI:** 10.1101/2021.02.23.432424

**Authors:** Christoph Gstöttner, Tao Zhang, Anja Resemann, Sophia Ruben, Stuart Pengelley, Detlev Suckau, Tim Welsink, Manfred Wuhrer, Elena Domínguez-Vega

## Abstract

As the SARS-CoV-2 pandemic is still ongoing and dramatically influences our life, the need for recombinant proteins for diagnostics, vaccine development, and research is very high. The spike (S) protein, and particularly its receptor binding domain (RBD), mediates the interaction with the ACE2 receptor on host cells and may be modulated by its structural features. Therefore, well characterized recombinant RBDs are essential. We have performed an in-depth structural and functional characterization of RBDs expressed in Chinese hamster ovary (CHO) and human embryonic kidney (HEK293) cells. To structurally characterize the native RBDs (comprising *N*- and *O*-glycans and additional posttranslational modifications) a multilevel mass spectrometric approach was employed. Released glycan and glycopeptide analysis were integrated with intact mass analysis, glycan-enzymatic dissection and top-down sequencing for comprehensive annotation of RBD proteoforms. The data showed distinct glycosylation for CHO- and HEK293-RBD with the latter exhibiting antenna fucosylation, higher level of sialylation and a combination of core 1 and core 2 type *O*-glycans. Additionally, from both putative *O*-glycosylation sites, we could confirm that *O*-glycosylation was exclusively present at T323, which was previously unknown. For both RBDs, the binding to SARS-CoV-2 antibodies of positive patients and affinity to ACE2 receptor was addressed showing comparable results. This work not only offers insights into RBD structural and functional features but also provides a workflow for characterization of new RBDs and batch-to-batch comparison.

## Introduction

Since the outbreak of COVID-19, the SARS-CoV-2 virus has infected more than 100 million individuals and influences our daily lives. The coronavirus is an enveloped RNA virus containing three different structural proteins in the membrane, the envelop (E) protein, the membrane (M) protein and the spike (S) glycoprotein [1]. The S protein is heavily glycosylated and forms a trimer on the SARS-CoV-2 surface. Each S protein carries 22 *N*-glycosylation sites [2] and consists of an S1 and an S2 subunit. Whereas the S2 subunit is necessary for membrane fusion, the S1 subunit directly interacts with the angiotensin converting enzyme 2 (ACE2) receptor in the human respiratory tract and facilitates the entry into the host cell [3]. In particular, the receptor binding domain (RBD) of the S1 subunit mediates the interaction with the ACE2 receptor [4]. This domain carries two *N*-linked glycans at positions N331 and N343 and, depending on the source, one or two *O*-linked glycosylation sites (T323/S325) are occupied [2, 5, 6]. The predicted *O*-glycosylation site at T323 is not present in the S protein of SARS-CoV-1 and has been hypothesized to have an influence on the interaction with the ACE2 receptor [7] and may be critical for conformational changes of the RBD [8]. Further studies have shown the role of glycosylation on ACE2-RBD binding. Crystallographic structures have confirmed the interaction between RBD and ACE2 receptor and highlighted the importance of ACE2 glycosylation in the interaction [9]. However, the RBD used was expressed in insect cells, and the role of RBD glycosylation was not addressed [9]. In a more recent publication the interaction of the *N*-glycan at position N343 of the RBD with the ACE2 receptor has been suggested by modelling the interaction using atomistic molecular dynamic simulations [10]. Another study revealed that the infectivity decreased by 1200 times (decrease of relative infectivity to 0.083%) when both *N*-glycosylation sites (N331 and N343) of the spike protein were silenced compared to the wild type versions. These findings suggest that glycosylation is involved in the binding to the receptor – either directly or by providing conformational stabilization [11]. In line with its role in virus-ACE2 interaction, the RBD is the primary target of neutralizing antibodies [12]. Interestingly, an antibody named S309, pulled from serum from a SARS-CoV-1 recovered patient, bound an epitope containing also glycosylation [13]. This binding site is also conserved in SARS-CoV-2 and was predicted as an epitope for neutralizing antibodies [14]. The neutralizing S309 antibody was shown to bind to a protein epitope (331-344) in combination with the glycan at N343. The antibody showed especially strong interaction with the core fucose and to a minor extent with the remaining glycan structure [13]. All these findings emphasize that the assessment of RBD glycosylation is highly important.

Recombinantly produced S proteins are essential tools in the fight against SARS-CoV-2, contributing to a further understanding of the interaction mechanism, providing efficient components for diagnostic purposes, and helping in vaccine development [15]. However, it is important to understand that glycosylation and other structural characteristics may differ considerably between different biotechnologically produced proteins and their natural forms. Considering the relevance of RBD glycosylation on ACE2 binding and recognition by neutralizing antibodies, the use of well-characterized S proteins is essential. The S protein, in particular, has been produced as full-length protein as well as in a short version containing the RBD [16]. Site-specific glycosylation analysis of the 22 *N*-glycosylation sites of the recombinant S glycoprotein expressed in human embryonic kidney 293 (HEK293) cells showed mainly complex type glycans but also, at certain glycosylation sites, high mannose structures and in lower amounts some hybrid structures [2, 5]. In particular, the two *N*-glycosylation sites (N331 and N343), which are located in the RBD were found to carry mainly complex type glycans as well as spurious amounts of high mannose glycans. Similarly, for the S1 subunit recombinantly produced in HEK293 cells, mainly complex type glycans were found at both sites [5]. As predicted by Uslupehlivan *et. al* [7] an *O*-glycosylation at the position T323 and/or at S325 was found. Whereas one report, analysing the whole S protein recombinantly produced, showed only trace levels of *O*-glycans [2] another study detected high levels of *O*-glycosylation [6]. Still, in these studies the localization of the *O*-glycosylation site at T323 and/or S325 was not possible. *O*-glycosylation at these two positions is also thought to be important to stabilize the conformation of the RBD or to introduce conformational changes [8]. All these studies have been performed using recombinant versions of the spike protein or subunits thereof. Interestingly, upon expression in HEK293 cells the amount of sialic acids varied from low to high depending on whether the complete S protein or only the S1 subunit was produced [5], suggesting that the *N*- and *O*-glycosylation is dependent on the context of the protein (S, S1 or RBD only). So far only terminal glycan epitopes of HEK293 produced RBD have been studied by NMR, however an assessment of the entire *N*-glycan structure, composition and the relative quantification of the *N*-glycans could not be achieved [17]. Furthermore, the *O*-glycosylation was neglected completely. Also, a characterization of the intact RBD is still missing, providing information on the combination of the glycans as well as on additional protein backbone modifications.

Here we present an in-depth structural and functional characterization of two commercially available SARS-CoV-2 RBDs produced in two different expression systems, HEK293 and Chinese hamster ovary (CHO) cells. To achieve comprehensive structural information, a multilevel characterization was performed (**Figure 1**). Both RBD samples were initially analyzed at the intact level by top-down sequencing using MALDI in-source-decay (MALDI-ISD) mass spectrometry (MS) [18, 19] and by sheathless capillary electrophoresis (CE)-MS after full *N*- and *O*-deglycosylation to establish their protein sequences and putative non-glycan modifications. With the established sequences and the help of sequential glycosidase treatment, the glycoforms on both *N*- and the *O*-glycosylation sites were assigned. Our findings were confirmed by glycopeptide analysis and the analysis of released *O*-glycans by porous graphitized carbon (PGC) nano-LC-ESI-MS/MS. To assess functional differences between the two RBD samples, we determined their binding characteristics to ACE2 and sera of patients who recovered from a previous SARS-CoV-2 infection.

**Figure 1:**
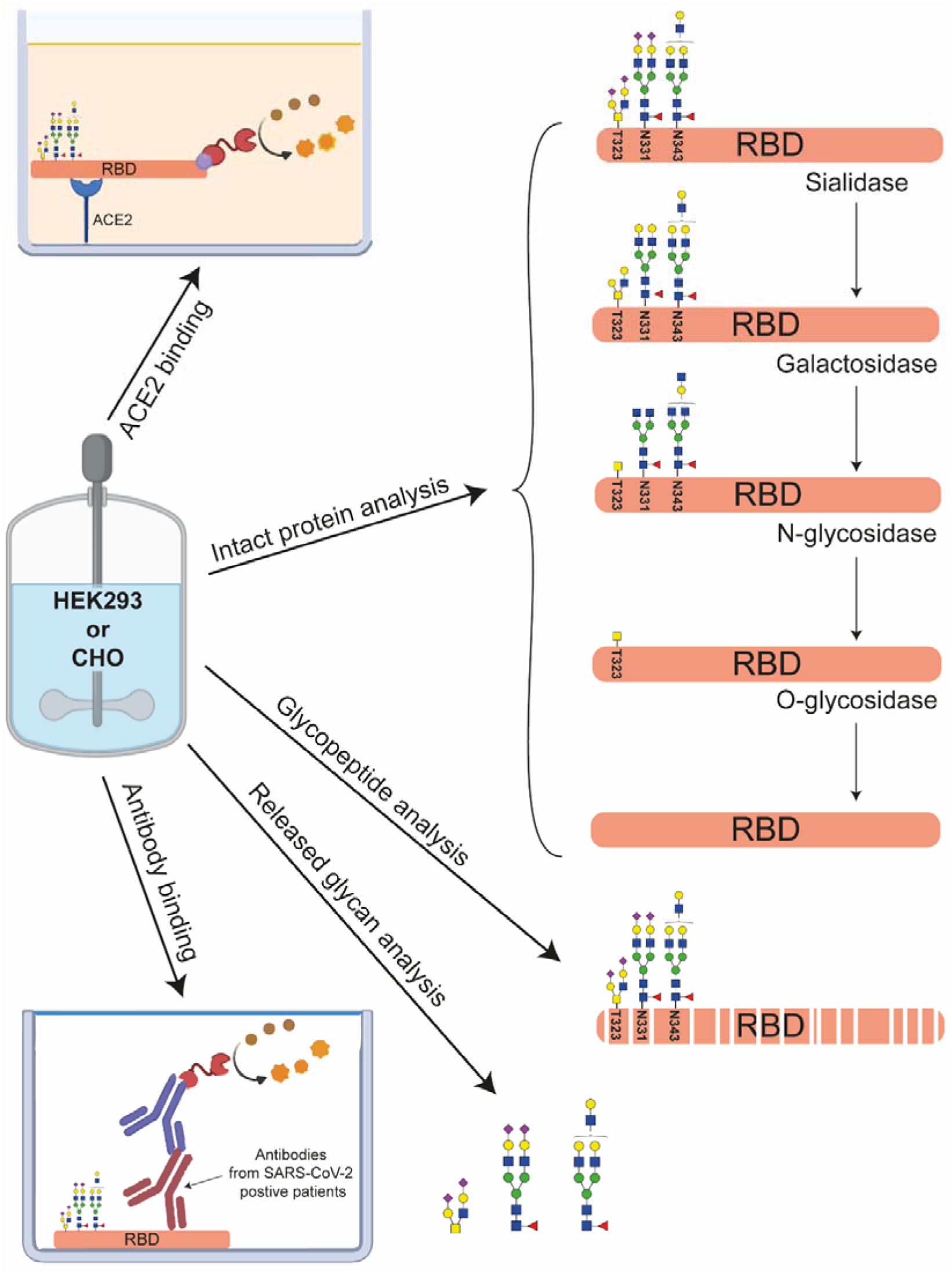
Multilevel characterization of RBDs produced in HEK293 or CHO cells. Next to intact protein analysis using CE-MS and MALDI-ISD-MS the samples are structurally characterized by glycan dissection, glycopeptide analysis and released glycan analysis. Their functional characterization was performed by measuring their binding characteristics to ACE2 and anti-SARS-CoV-2 antibodies.

## Materials and Methods

### Reagents and samples

Ethanol (absolute for analysis) and 2-propanol (for analysis) were obtained from Merck (Darmstadt, Germany). Sodium chloride (≥99.8%), 7.5 M ammonium acetate solution (for molecular biology), Tris(hydroxymethyl)aminomethane (≥99.8%), sodium hydroxide (≥99.8%), methanol anhydrous (≥99.8%), hydrogen chloride (ACR reagent grade 37%), ammonium bicarbonate (≥99.0%), dithiothreitol (DTT) (≥98.0%), iodoacetamide (IAA) (≥99.5%), trifluoroacetic acid (TFA) (≥99.0%), Dowex cation-exchange resin (50W-X8) and sodium borohydride (≥98.0%) was obtained from Sigma Aldrich (St. Louis, Missouri). Potassium hydroxide (≥85.0%) was obtained from Honeywell Fluka (Charlotte, NC). Acetic acid (≥99.7%) was purchased from VWR (Radnor, PA). Methanol (Ultra LC-MS grade) was purchased Actu-All Chemicals (Oss, The Netherlands). 8 M guanidine hydrochloride (GuHCl) (for protein biology) was obtained from Thermo Fisher Scientific (Waltham, MA). HPLC SupraGradient acetonitrile was obtained from Biosolve (Valkenswaard, The Netherlands). 50% trimethoxysilylpropyl modified polyethylenimine (PEI) dissolved in 2-propanol was provided by Gelest (Morrisville, NC). Ultrapure water was used for the all preparations and washes, generated from a Q-Gard 2 system (Millipore). Tetramethylbenzidine was obtained from Surmodics (Eden Prairie, MN).

Recombinant RBDs (Wuhan-Hu-1-isolate (MN908947)), either transiently expressed in HEK293 or stably expressed in CHO cells, were used (InVivo Biotech Services, Henningsdorf, Germany). The constructs contained the amino acid sequence 319 to 541 with a C-terminal 6xHis-Tag. Recombinant RBDs were purified using immobilized metal affinity chromatography and a size exclusion polishing step. The samples were stored in 20 mM sodium phosphate, 300 mM NaCl, pH 7.2. Glycosidases SialEXO (sialidases α2-3, α2-6 & α2-8), GalactEXO (galactosidases β1-3 and β1-4), OglyZOR (endo-α-N-acetylgalactosaminidase), OpeRATOR (*O*protease), α1-2 fucosidase and α1-3,4 fucosidase were obtained from Genovis (Lund, Sweden). Peptide *N*-glycosidase F (PNGaseF) was purchased from Roche Diagnostics (Mannheim, Germany). Trypsin and elastase were purchased from Promega (Madison, WI). Super-DHB (sDHB) and a 384 MTP BigAnchor target plate (Bruker Daltonics, Bremen, Germany) were used for MALDI-ISD analysis. The calibrant ubiquitin was purchased from Sigma-Aldrich, Germany.

### Intact RBD analysis by sheathless CE-MS

Samples for intact protein analysis were buffer exchanged with Tris pH 6.8 using 10□kDa Vivaspin MWCO filters Sartorius (Göttingen, Germany) to a final concentration of 1 μg/μL. Subsequently the enzymes SialEXO, GalactEXO, OglyZOR or OpeRATOR were added according to the producers specifications and the mixtures were incubated overnight at 37°C. For samples treated with PNGaseF alone or in combination with other glycosidases, PNGaseF was added in a ratio 1:5 (v/v) and incubated overnight at 37°C. For removing antenna fucosylation 20 μg of HEK293-RBD sample was incubated with 4.4 μg fucosidase α1-2 or 8.8 μg fucosidase α1-3,4 overnight at 37°C. Afterwards all samples were buffer exchanged with 100 mM ammonium acetate pH 3.0 and subsequently analyzed by sheathless CE-MS.

For sheathless CE measurements a Sciex CESI 8000 instrument combined with a Bruker Impact qTOF-MS was used. Bare-fused silica CE capillaries equipped with a porous tip were obtained from Sciex (Framingham, MA). To avoid protein adsorption fused silica capillaries were permanently positively coated with PEI following the protocol of Sciex [20]. To assess the performance of the PEI coated capillary, a protein test mixture was analyzed. For intact RBD analysis a background electrolyte (BGE) consisting of 9% acetic acid and 10% methanol was used. The separation capillary and the conductive line were filled with the BGE for 3 min (100 psi forward pressure) and 1 min (75 psi reverse pressure), respectively. Afterwards the sample was injected for 15 s with a pressure of 5 psi (11.2 nL; 1.75% of the total capillary volume) followed by a plug of BGE for 25 s with a pressure of 0.5 psi. The separation was carried out by applying 20 kV (reversed polarity) at 20°C for 45 min. After the separation was completed the voltage was ramped down to 1 kV in 5 min with a pressure of 50 psi (forward and reverse).

The CE was coupled to MS using a nano-electrospray ionization source. The system was operated in positive ion mode with a dry gas flow rate of 2.0 L/min, a dry temperature of 120°C and a capillary voltage of 1300 V. The collision cell voltage was set to 40 eV and the quadrupole energy to 5 eV. An in-source collision-induced dissociation energy (applied between funnel 1 and 2) of 50 eV was used. The transfer time and the pre-pulse storage time were set to 150 μs and 20 μs, respectively. The monitored *m/z* range was 500-4000.

### ESI Data analysis of intact RBD

Intact RBD data were deconvoluted using the DataAnalysis 5.3 (Bruker Daltonics) maximum entropy algorithm. Deconvoluted mass spectra were baseline subtracted and smoothed using the Gaussian smoothing function with a width of 0.5 Da and one cycle. Glycoforms were assigned based on the average masses observed and observed mass shifts with enzyme treatments. Additionally, data obtained from *N*-glycopeptide analysis and *O*-glycan release was used to confirm findings and distinguish glycan isomers.

### MALDI-ISD top-down protein sequence analysis

Fully deglycosylated RBD samples were reduced for 30 min at 50°C using DTT. Two μL of the reduced sample were spotted on a hydrophilic anchor of an MTP BigAnchor sample plate and incubated. After 2 min, the remaining droplet was removed and the spot was washed using 0.1% TFA in water. Subsequently, 1 μL of sDHB matrix solution (25 μg/μL in 50% acetonitrile/49.9% water/ 0.1% TFA) was deposited on the dried sample spot. MALDI-ISD spectra were acquired with a rapifleX MALDI-TOF MS instrument (Bruker) in positive reflector ion mode using a method optimized for ISD acquisition provided by the manufacturer. The detection voltage was set to 3.26 kV and a sum of 35,000 laser shots were accumulated at 1000 Hz at elevated laser power adapted to the sDHB matrix. The spectra were processed in FlexAnalysis (Compass for flexSeries 2.0) using a smoothing step (5 cycles 0.15 Da) and baseline subtraction (TopHat). Peaks were picked using the SNAP2 algorithm up to 5,000 Da and the SNAP algorithm up to 10,000 Da. The spectra were calibrated using a MALDI-ISD spectrum from 50 pmol bovine ubiquitin prepared with sDHB. For interpretation and annotation of ISD spectra BioTools 3.2 SR7 was used (Bruker).

### Glycopeptide analysis by RPLC-MS/MS

For glycopeptide generation a double digestion with trypsin followed by elastase was performed. Here, 300 μg (259 μL) of RBD protein was reduced with 10 μL DTT (45 mM) for 30 min at 50°C. Afterwards the protein was alkylated with 5 μL IAA (100 mM) for 30 min at 37°C in the dark followed by addition of 10 μL DTT (45 mM) and incubation for 15 min at 50°C to stop the alkylation reaction. The samples were allowed to cool down to 37°C and incubated with 60 μL Trypsin (100 ng/μL) for 12 h at 37°C. To stop the digestion reaction the samples were heated for 1 min to 90°C and the concentration was determined using a NanoDrop spectrophotometer (Thermo Fisher Scientific) resulting in a protein concentration of 0.94 μg/μL. For elastase digestion 120 μg (127 μL) of tryptic digested RBD was incubated with 2.4 μL (1 μg/μL) elastase for 12 h at 37°C. The digestion reaction was stopped by adding 1.3 μL of formic acid. DTT, IAA, trypsin and elastase were dissolved in 50 mM ammonium bicarbonate buffer with a pH of 7.8.

The digested samples were separated using a nanoElute (Bruker) nanoflow UHPLC equipped with an aurora 25 cm x 75 μm C18 column with a particle size of 1.6 μm (IonOpticks, Parkville, Victoria, Canada). For each run 800 ng of digested protein was injected. The separation was performed using water and acetonitrile as mobile phase A and B, each containing 0.1% formic acid. A linear gradient from 2 to 35% in 90 min was used for analyte separation with a column temperature of 50°C. The nano-LC was connected to a timsTOF Pro (Bruker) using a nanoBooster (Bruker) supplying acetonitrile enriched dopant gas (0.2 bar). The capillary voltage was set to 1400 V and a dry gas of 3.0 L/min with a temperature of 180°C was used. The quadrupole ion energy was set to 5 eV and the collision cell collision energy to 7 eV. A pre-pulse storage time of 10 μs was used. AutoMS/MS data was recorded as follows: for MS measurements a *m/z* range between 600 and 3000 was monitored and spectra were acquired with a speed of 2 Hz; for MS/MS measurements a *m/z* range of 50 to 3000 was monitored and a dynamic spectra acquisition between 4 and 10 Hz was used depending on target intensity, which was set to 8000 counts. For fragmentation preferably charge states between 3+ and 6+ were selected. The collision energy ramp settings were adapted from Hinneburg *et al*. [21] with a collision energy ramp from 31 eV at *m/z* 700 to 80 eV at *m/z* 1500. The collision energy stepping was 100% for 40% of the MS/MS spectra (preferred fragmentation of peptide moiety) and 50% for 60% of the MS/MS spectra (preferred fragmentation of glycan moiety).

### Data analysis of glycopeptides

MS/MS data of N-linked glycopeptides were processed using DataAnalysis 5.3 and MGF peak lists were imported into BioPharma Compass 2021 (Bruker) and further analyzed using the glycopeptide analysis workflow. This workflow includes MS/MS spectra classification using typical fragmentation patterns to determine the peptide mass and the glycan mass of a glycopeptide. The peptide sequences were then identified using the theoretical digest feature of the software. In a second search, the classified MS/MS spectra were submitted to a glycan search using the GlycoQuest search engine within BioPharma Compass 2021 software. The most intense isotope of each charge state ([M+H]^+^, [M+2H]^2+^, or [M+3H]^3+^) was extracted and the glycopeptides were quantified and results normalized to the total peak intensity of all glycopeptides within one sample to 100%.

### Released *O*-glycan analysis by PGC nano-LC-ESI-MS/MS

*O*-glycan alditols released from RBD samples were prepared using a 96-well plate sample preparation method as previously described [22]. In brief, 20 μg of RBD samples were applied to the hydrophobic Immobilon-P PVDF membrane (Millipore, Amsterdam, The Netherlands) in a 96-well plate format. Protein denaturation was achieved by addition of 75 μL denaturation mix (72.5 μL 8 M GuHCl and 2.5 μL 200 mM DTT) to each well, followed by shaking for 15 min and incubation at 60°C in a moisture box for 30 min. Subsequently the unbound material was removed.

The *N*-glycans were released by addition of PNGaseF (2 U of enzyme diluted with water to 15 μL) to each well and overnight incubation at 37 °C. Released *N*-glycans were removed from the PVDF plate by centrifugation. After removal of *N*-glycans, the *O*-glycans were released from the PVDF membrane immobilized sample via reductive β-elimination. Briefly, 50 μL of 0.5 M NaBH_4_ in 50 mM KOH was applied onto each PVDF membrane well after rewetting with 3 μL of methanol. Plates were shaken for 15 min on a horizontal shaker and incubated in a humidified plastic box for 16 h at 50°C. After incubation and cooling to RT, released *O*-glycans were recovered by centrifugation at 1000 g for 2 min into 96-well collection plates. The wells were rewetted by 3 μL of methanol and washed three times with 50 μL of water with 10 min incubation steps on a horizontal shaker prior to centrifugation at 500 g for 2 min. Before desalting the collected samples were concentrated to approximately 30 μL under vacuum in a SpeedVac concentrator at 35°C for 2 h. Subsequently, 3 μL of glacial acetic acid were added to quench the reaction followed by brief centrifugation to collect the sample at the bottom of the well. Collected *O*-glycan alditols were then desalted and purified followed by a desalting and PGC clean-up using a 96-well plate based protocol [22]. Samples were dried in a SpeedVac concentrator directly in polymerase chain reaction (PCR) plates and redissolved in 10 μL of water prior to PGC nano-LC-ESI-MS/MS analysis.

The analysis of *O*-glycan alditols was performed following a method described previously [22]. Measurements were performed on an Ultimate 3000 UHPLC system (Dionex/Thermo) equipped with an in house-packed PGC trap column (5 μm Hypercarb, 320 μm x 30 mm) and an in-house-packed PGC nano-column (Grace Discovery Sciences, Columbia, MD, USA) (3 μm Hypercarb 100 μm x 150 mm) coupled to an amaZon ETD speed ion trap (Bruker). Mobile phase A consisted of 10 mM NH_4_HCO_3_, while mobile phase B was 60% (v/v) acetonitrile/10 mM NH_4_HCO_3_. To analyze glycans, 2 μL of sample prepared from 4 μg glycoprotein was injected and trapped on the trap column using a 6 μL/min loading flow of 1% buffer B for 5 min. Separation was achieved with a gradient of B: 1-52% over 72 min followed by a 10 min wash step using 95% of B at a flow of rate of 0.6 μL/min. The column was held at a constant temperature of 45°C. Ionization was achieved by using 2-propanol enriched nitrogen gas at 3 psi provided by a nanoBooster source (Bruker). The capillary voltage was set to 1000 V and a dry gas temperature of 280°C at 5 L/min was used. MS spectra were acquired within an *m/z* range of 380-1850 using negative ion mode. Smart parameter setting (SPS) was set to *m/z* 900. MS/MS spectra were recorded using the top three highest intensity peaks.

### Data analysis of released *O*-glycans

Structures of detected glycans were confirmed by MS/MS in negative mode [23]. Glycan structures were assigned on the basis of the known MS/MS fragmentation patterns in negative-ion mode [24–26], elution order, and general glycobiological knowledge, with help of Glycoworkbench [27] and Glycomod software [28]. Relative quantification of individual glycans was performed by normalizing the total peak area of all glycans within one sample to 100%. Relative abundances of specific glycan structures were grouped by summing relative abundances of each glycan multiplied by the number of motifs per glycan.

### SARS-CoV-2-IgG ELISA with RBD antigens

The antigens (intact or glycosidase treated HEK293-RBD or CHO-RBD) were used to coat immunoassay plates at a concentration of 2 μg/ml. Subsequently, blocking was performed with blocking buffer containing 1% bovine serum albumin (BSA). Serum collected >10 weeks after onset of first symptoms from 12 SARS-CoV-2 PCR positive tested donors was applied to the RBD-coated wells at a dilution of 1:101. As a negative control, a 1:101 diluted serum pool containing 10 sera taken from healthy individuals in the year 2012 was used. Subsequently, bound IgG from sera was detected by anti-human IgG-HRP (Seramun Diagnostica GmbH, Heidesee, Germany) and developed with tetramethylbenzidine. The reaction was stopped using sulfuric acid. The absorbance was measured at 450 nm using a microplate reader. In order to determine the linear correlation between the absorption values, the Pearson correlation was calculated using GraphPad Prism 9. Linear regression was employed for the visualization of the correlation. For this analysis, the negative control value was excluded.

### ACE2 receptor binding assay

Intact and glycosidase treated RBD, S1 subunit and the S protein (InVivo Biotech) were biotinylated using 10 molar excess of NHS-LC-Biotin (Thermo Scientific). Immunoassay plates were coated with 2.5 μg/ml recombinant angiotensin converting enzyme-2 (ACE2, Sigma Aldrich). Subsequently, the assay plate wells were blocked using blocking buffer containing 1% BSA. Dilutions of the biotinylated antigens ranging from 1 μg/ml to 0.001 μg/ml were applied to the ACE2-coated assay plate wells in duplicates. Bound biotinylated antigens were detected using streptavidin peroxidase conjugate (Roche) and developed with TMB. The reaction was stopped using sulfuric acid. The absorbance was measured at 450 nm using a microplate reader. GraphPad Prism 9 was used to plot log(dose) response curves (variable slope, four parameters) and to compute nonlinear fits which were utilized to calculate the half-maximal concentrations (EC_50_).

## 3. Results and Discussion

### 3.1. Structural characterization of CHO and HEK293 produced RBDs

We characterized the RBD domains (amino acids 319-541 containing a C-terminal His-tag) produced in CHO and HEK293 cells. This domain contains two *N*-glycosylation sites at positions N331 and N343 as well as two potential *O*-glycosylation sites at T323 and S325. Analysis of the intact RBDs revealed a complex pattern of signals comprising different *N*- and *O*-glycans and additional protein backbone modifications. Therefore, to unravel this heterogeneity and achieve comprehensive structural characterization, we used a multilevel approach (**Figure 1**). Next to classical released glycan and glycopeptide approaches, we applied a step-by-step dissection of glycans at the intact level similar to the work of Wohlschlager *et. al* for the biopharmaceutical etanercept [29].

#### 3.1.1. Characterization of the protein backbone after *N*- and *O*-glycan removal

To get information on the integrity of the protein backbone, the RBD proteins were treated with PNGaseF to remove the *N*-glycans and with an endo-α-N-acetylgalactosaminidase in combination with a mix of sialidases to remove the *O*-glycans. After deglycosylation, the RBDs were analyzed by CE-MS for intact mass and MALDI-ISD MS for top-down sequencing.

Analysis of the samples by CE-MS resulted in one main peak in the base peak electropherogram (BPE) corresponding to the RBD. For the RBD produced in CHO cells, the observed averaged mass (26033.9 Da) was 119.0 Da higher than the theoretical mass calculated solely based on the amino acid sequence (25914.9 Da) (**Figure 2A**). As the RBD contains a free cysteine at position C538 [30], it was presumably cysteinylated (+119.1 Da). This modification is often observed for free light chains during antibody production using CHO cells [31]. After reducing the disulfide bonds with DTT, the mass of the deglycosylated and completely reduced protein was 25923.0 Da, which is consistent with the expected theoretical mass (25923.0 Da) confirming the presence of cysteinylation (data not shown). For the RBD expressed in HEK293 cells, a deconvoluted mass of 26144.9 Da was observed (**Figure 2B**). Considering a cysteinylation of the free cysteine also in the HEK293-RBD, an additional mass difference of 110.9 Da was observed compared to the theoretical mass indicating additional modifications.

**Figure 2:**
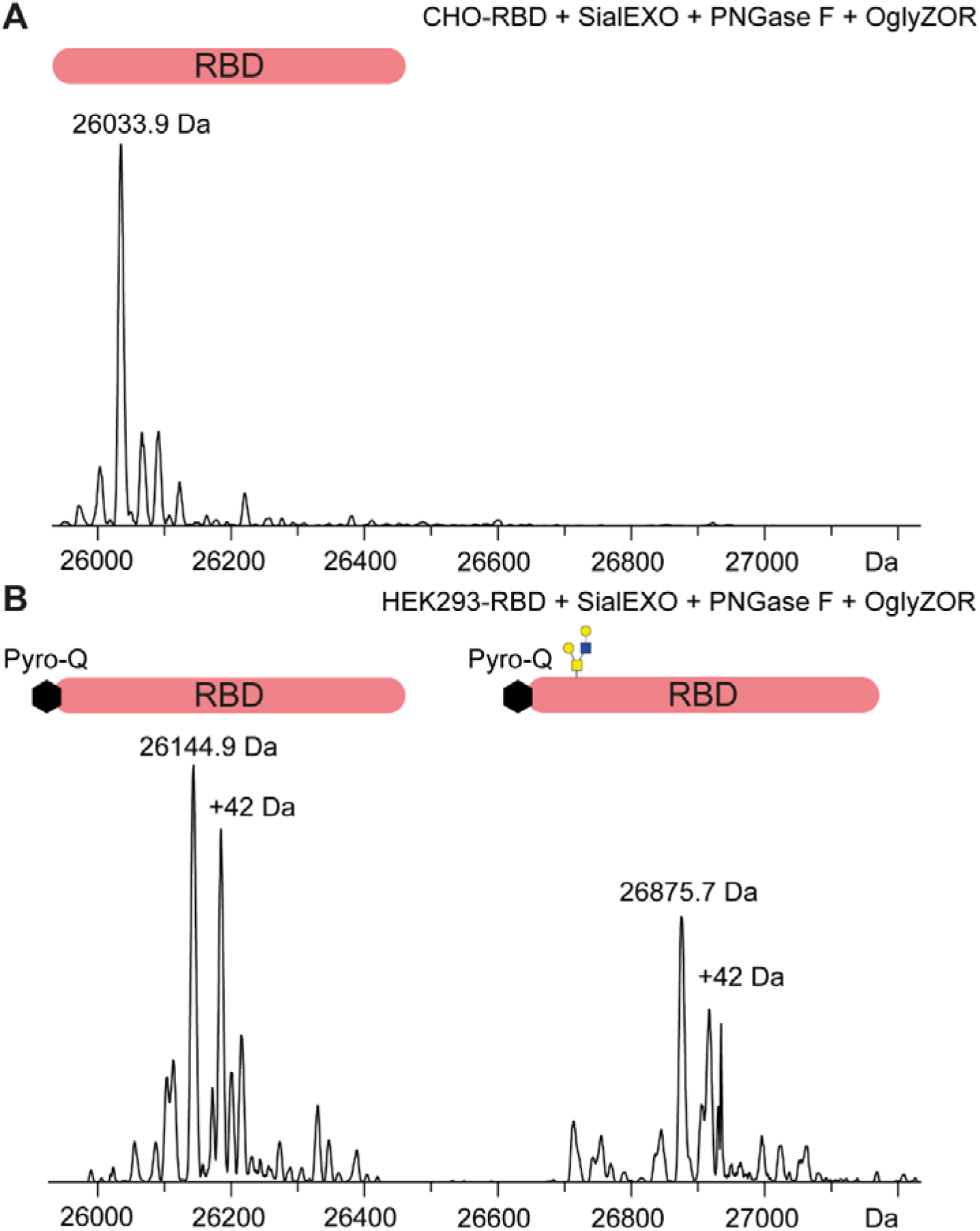
Deconvoluted ESI mass spectra of the main peak observed after sheathless CE-MS for A) CHO-RBD and B) HEK293-RBD. HEK293-RBD carries an additional N-terminal pyroglutamate. Blue square, N-acetylglucosamine; yellow square, N-acetylgalactosamine; yellow circle, galactose.

Both RBD samples were analyzed at the intact level by top-down sequencing using MALDI-ISD MS after full reduction and *N*- and *O*-deglycosylation. MALDI-ISD provides large terminal sequence tags and detects terminal (and internal) PTMs from intact proteins. For both RBDs we detected a C-terminal sequence tag of 73 amino acids (RMS error of 0.028 Da) verifying the expected C-terminus of the sequence. The N-terminal sequence tag, however, was not consistent between both RBDs. While the CHO-RBD sequence was in full agreement with the MALDI-ISD spectrum (data not shown), the HEK293-RBD sequence was found to be N-terminally extended by pyro-Glu (mass shift of 111.03 Da). 52% of the sequence were confirmed this way including an N-terminal sequence tag of 46 residues for the nonglycosylated and 29 residues for the *O*-glycosylated form (Supplementary **Figure S1**). Of note, the HEK293-RBD came with a slightly different sequence compared to the CHO-RBD, resulting in an additional glutamine at the N-terminus after cleavage of the signaling peptide (**Figure S1**). Taking pyroglutamic acid formation into consideration, the measured mass perfectly fits the theoretical mass (26145.1 Da). The N-terminal pyroglutamic acid formation was further confirmed by bottom-up analysis (data not shown). In addition, we found a species migrating before the main signal in the BPE with a +42.3 Da mass difference (26187.4 Da). This difference in mass might correspond to acetylation, however, it could not be confirmed by bottom-up analysis after trypsin digestion. This species was only observed in HEK293-RBD and not in CHO-RBD.

#### 3.1.2. RBD *O*-glycan characterization

The RBD has two potential *O*-glycosylation sites at position T323 and S325. To characterize the *O*-glycans, the de-*N*-glycosylated RBD (after treatment with PNGaseF) and the released *O*-glycans were analyzed by CE-MS (**Figure S2**) and PGC nano-LC-ESI-MS/MS (**Figure 3**), respectively. **Table S1** summarizes the obtained results. In CHO-RBD mainly core 1 structures with 2 sialic acids (H1N1S2) were observed. HEK293-RBD showed a much more diverse *O*-glycosylation pattern with a core 1 structure with 2 sialic acids as the main signal. Additionally, several core 2 structures, with and without fucose as well as with sulfation were detected. These data are in accordance with the data in **Figure 2B** for the sample treated with an endo-α-N-acetylgalactosaminidase, where an additional signal of 26875.7 Da was observed. This mass corresponds to an H2N2 modification (theoretical mass 26875.8 Da) which is presumably a core 2 *O*-glycan which cannot be cleaved by the endo-α-N-acetylgalactosaminidase. Similar glycoforms were not detected in CHO-RBD. The position of fucoses, either to the terminal galactose or the *N*-acetylglucosamine were confirmed by MS fragmentation and treatment with different fucosidases (**Figure S3**).

**Figure 3:**
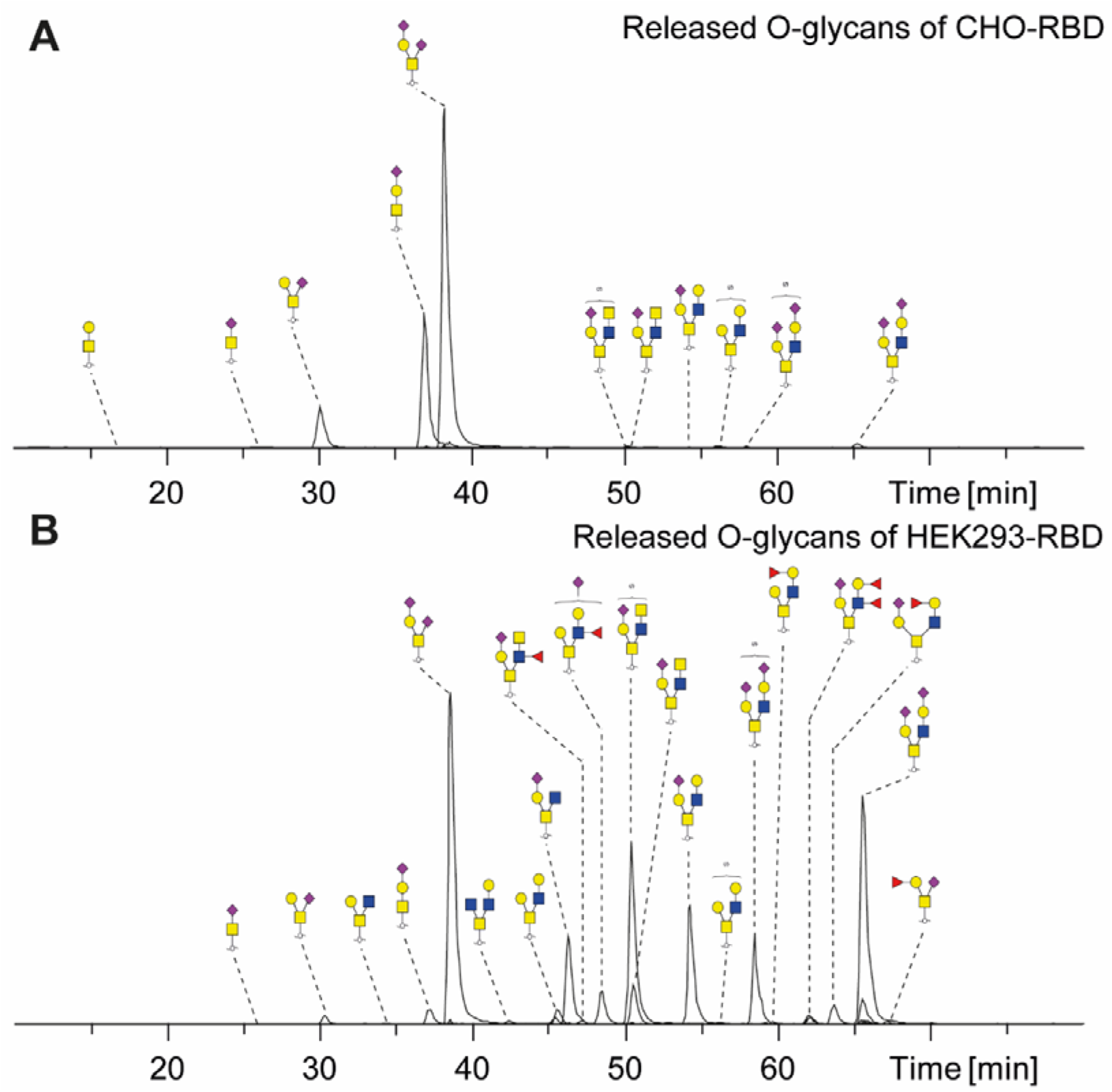
PGC nano-LC-ESI-MS/MS of released *O*-glycans from A) CHO-RBD and B) HEK293-RBD. Yellow square, N-acetylgalactosamine; yellow circle, galactose; blue square, N-acetylglucosamine; red triangle, fucose; purple diamond, N-acetylneuraminic acid (sialic acid), S, sulfate.

Hitherto published S protein bottom-up studies have not revealed the *O*-glycan attachment site, leaving both T323 and S325 as valid options [2, 6]. To resolve this, we follow a different strategy based on *N*-glycan removal (with PNGaseF) and *O*-protease cleavage using the enzyme OpeRATOR in combination with a mix of sialidases. The *O*-protease cleaves the protein at the N-terminal site of an *O*-glycosylation site (**Figure S3**). This would result in a loss of the amino acids 319-322 from the RBD if the *O*-glycosylation is on T323 or 319-324 in case S325 carries the *O*-glycan. As shown in **Figure S4A,** only one signal with a mass of 25919.0 Da was observed for the CHO material, which correlates to the RBD cleaved at T323 with a core 1 *O*-glycan H1N1 (theoretical mass 25918.8 Da). No RBD cleaved at S325 was observed. In HEK293-RBD, the same signal with a core 1 glycan H1N1 was observed also indicating that the glycosylation is located at T323 (**Figure S4B**). Next to this signal, an amount of uncleaved RBD with presumably core 2 *O*-glycan structures was detected. This is in line with a recent article that shows that the used *O*-protease can cleave N-terminal to core 1 but not core 2 glycan structures, similar to endo-α-N-acetylgalactosaminidase [32]. To confirm that the core 2 structures are located at T323 we performed MALDI-ISD MS analysis using super-DHB as a matrix. In contrast to CID fragmentation in bottom-up analysis, MALDI-ISD is known to result in mainly singly charged c- and z+2 ions with labile modifications remaining intact, which allows to localize *O*-linked glycosylation sites. MALDI-ISD MS of CHO-RBD after *N*-glycan and sialic acid removal showed N-terminal fragments (c-ions) with core 1 structure (H1N1) for the site T323 and no additional glycosylated fragments at S325 (+365.1 Da) could be detected confirming the presence of the *O*-glycosylation at T323 (**Figure S5A**). No fragments of T323 without glycosylation were observed indicating full site occupancy. The analysis of HEK293-RBD comprising only core 2 structure *O*-glycans (after *N*-glycan and core 1 *O*-glycan removal) clearly showed that the core 2 structures are also located on T323 (**Figure S5B**). In conclusion, the combination of intact analysis of *O*-protease treated RBDs together with MALDI-ISD allowed location of the *O*-glycans to the T323 with high confidence.

#### 3.1.3 Assessment of RBD *N*-Glycosylation

We studied the two *N*-glycosylation sites N331 and N343 using a bottom-up glycopeptide approach combined with a step-by-step dissection of glycans at the intact level [29]. Whereas the glycopeptide data yielded the glycan composition per site, we studied the combination of these glycans at the intact level [33].

The intact RBDs were first incubated with a mixture of sialidases and galactosidases. These enzymes removed the sialic acids and the terminal galactoses on the *N*- as well as *O*-glycans, resulting in considerably simplified deconvoluted mass spectra as shown in **Figure S6** and **Figure S7**. Overall, mainly fucosylated complex type glycans were observed for both RBDs (>90.0%). This is in line with previous studies on HEK293 produced intact S protein or S1 subunit [2, 5]. The dissection of the terminal galactoses also permits direct distinction between *N*-acetyllactosamine (LacNAc) repeats and additional antenna. CHO-RBD showed a higher relative abundance of LacNAc repeats than HEK293-RBD. Additionally, CHO-RBD showed a higher antennarity, with two triantennary structures as the most abundant glycoforms, contrary to HEK293 material with di- and triantennary glycans as major signals. This was confirmed by the analysis of the glycopeptides (**Figure S8, Table S2 and S3**). In general, more di- and triantennary structures were observed for HEK293-RBD (N331: 78.4% and N343: 84.5%) compared to the CHO-RBD (N331: 48.1% and N343: 72.1%). Therefore, CHO-RBD showed a larger contribution of tetraantennary structures and LacNAc repeats (N331: 42.7% and N343: 24.4%) compared to HEK293-RBD (N331: 15.7% and N343: 9.9%). Between both glycosylation sites a difference in the number of LacNAc repeats and high-antennary structures was observed with minor amounts on N343 compared to N331.

In addition, we found spurious amounts of hybrid type glycans only at N343 and only in HEK293-RBD (0.8%), similar to previous publications [2, 5]. Regarding high mannose glycans, low amounts were detected for both RBDs. In the case of HEK293-RBD, Man5 and Man6 glycans with and without phosphorylation were observed, whereas in the case of CHO-RBD Man5 and Man6 glycans carrying an additional phosphate were detected. For HEK293-RBD the high mannose and phosphorylated high mannose structures show combined abundances of 3.1% (N331) and 1.3% (N343) in line with the findings of Watanabe *et al*. [2] for full-length S protein. For CHO-RBD the distribution was more skewed with 7.7% (N331) and 0.9% (N343).

Besides, in the deconvoluted mass spectrum of the HEK293-RBD treated with sialidase and galactosidase a pattern of signals with a mass shift of +308.1 Da was observed. A combination of one galactose and one fucose could explain this mass shift. It was shown in literature that β-galactosidases are not able to remove the terminal galactose if an antenna fucose, either linked to the galactose itself or to the *N*-acetylglucosamine, is present [34]. Therefore, we incubated the HEK293-RBD either with α1-2 fucosidase and in parallel with α1-3,4 fucosidase to remove fucoses linked to the galactose and *N*-acetylglucosamine, respectively. Incubation with these fucosidases allowed the β-galactosidase to remove the terminal galactose. As shown in **Figure S3,** after removing the differently linked antenna fucoses, the +308.1 signals disappear completely confirming antenna fucosylation and providing information on the linkage of the antenna fucose with a predominant linkage to the N-acetylglucosamine. Additionally, antenna fucosylation in HEK293-RBD was confirmed by glycopeptide analysis (26.7% and 33.8% of antenna fucosylation on N331 and N343, respectively) **(Figure S9, Table S2 and S3).** No antenna fucosylation was found for CHO-RBD with any of the approaches.

The analysis of the intact RBDs without any previous enzymatic treatment resulted in a very complex mass spectrum (**Figure 4**). Based on the information obtained for the released *O*-glycans, glycopeptide and enzymatically-treated intact RBDs the spectra were confidently assigned. In particular, the HEK293-RBD exhibited a large heterogeneity. In addition to the variability resulting from the acetylation and antenna fucosylation, which were not observed in CHO-RBD, also a higher degree of sialylation was observed for HEK293-RBD. This was also supported by the glycopeptide data in which the CHO-RBD showed only 2.5% or 0.8% sialylated glycans, whereas the HEK293-RBD contained 56.4% or 36.3% on N331 or N343, respectively (**Figure S10, Table S2 and S3**). Furthermore, in CHO-RBD only monosialylated species were observed, while in the HEK293-RBD mono-, di- or trisialylated species were detected. Interestingly, these high sialylation levels were not observed by Watanabe *et al*. who reported only 22% sialylation on N331 and 4% sialylation on N343 for the full length S protein [2]. This might be due to the different constructs with RBD expression versus S1 subunit or S-protein. A similar effect of lower sialylation on the complete S protein has also been previously reported by comparing the entire S protein expressed in HEK293 cells with the S1 subunit [5]. Whereas S protein carried either ~ 40% or 10% sialylation on N331 or N343, the S1 subunit carried 80% or 50% on N331 or N343, respectively. These findings highlight that sialylation, which influences the isoelectric point of a protein, can change with the length of protein expressed (S, S1 or RBD) and must be taken into account.

**Figure 4:**
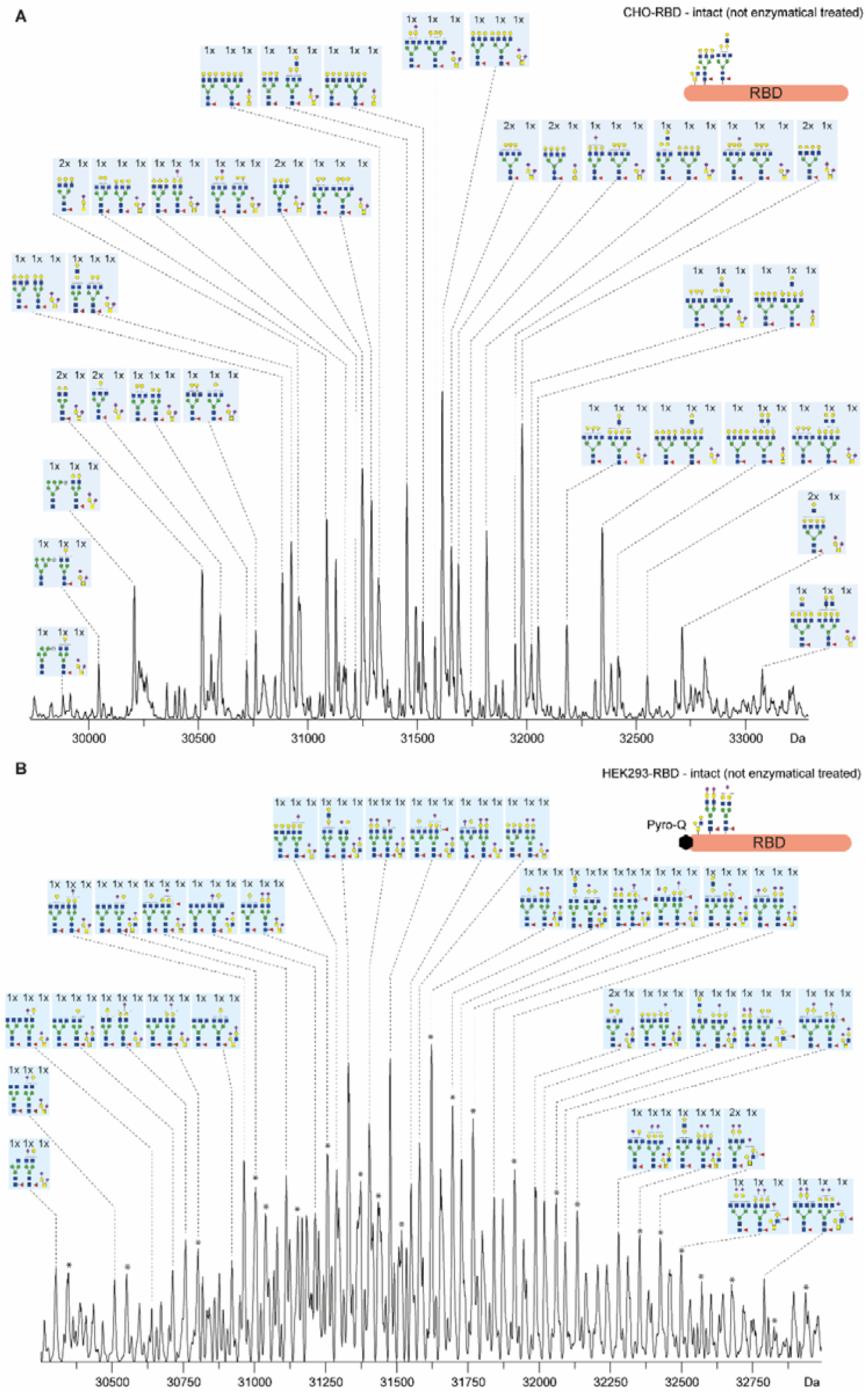
Deconvoluted mass spectra of the intact RBD (not enzymatically treated) produced by A) CHO cells or B) HEK293 cells. The assignments were based on previous enzyme treatments, mass and glycopeptide as well as released O-glycan data. Peaks marked with an asterisk * are presumably the acetylated variant of RBD. Yellow square, N-acetylgalactosamine; yellow circle, galactose; blue square, N-acetylglucosamine; red triangle, fucose; purple diamond, N-acetylneuraminic acid (sialic acid).

Finally, to support our assignments, the intact mass spectra were reconstructed from the glycopeptide data as described [33]. **Figure 5A**, shows a very strong correlation for the CHO-RBD confirming the assignments. For HEK293-RBD reconstructed mass spectrum showed a shift towards lower masses compared to the intact profile (**Figure 5B**). Proteins with higher sialylation levels often show a discrepancy between the intact and the glycopeptide-reconstructed mass spectra [33] which could be attributable to a non-random combination or to a biased ionization efficiency of sialylated glycopeptides.

**Figure 5:**
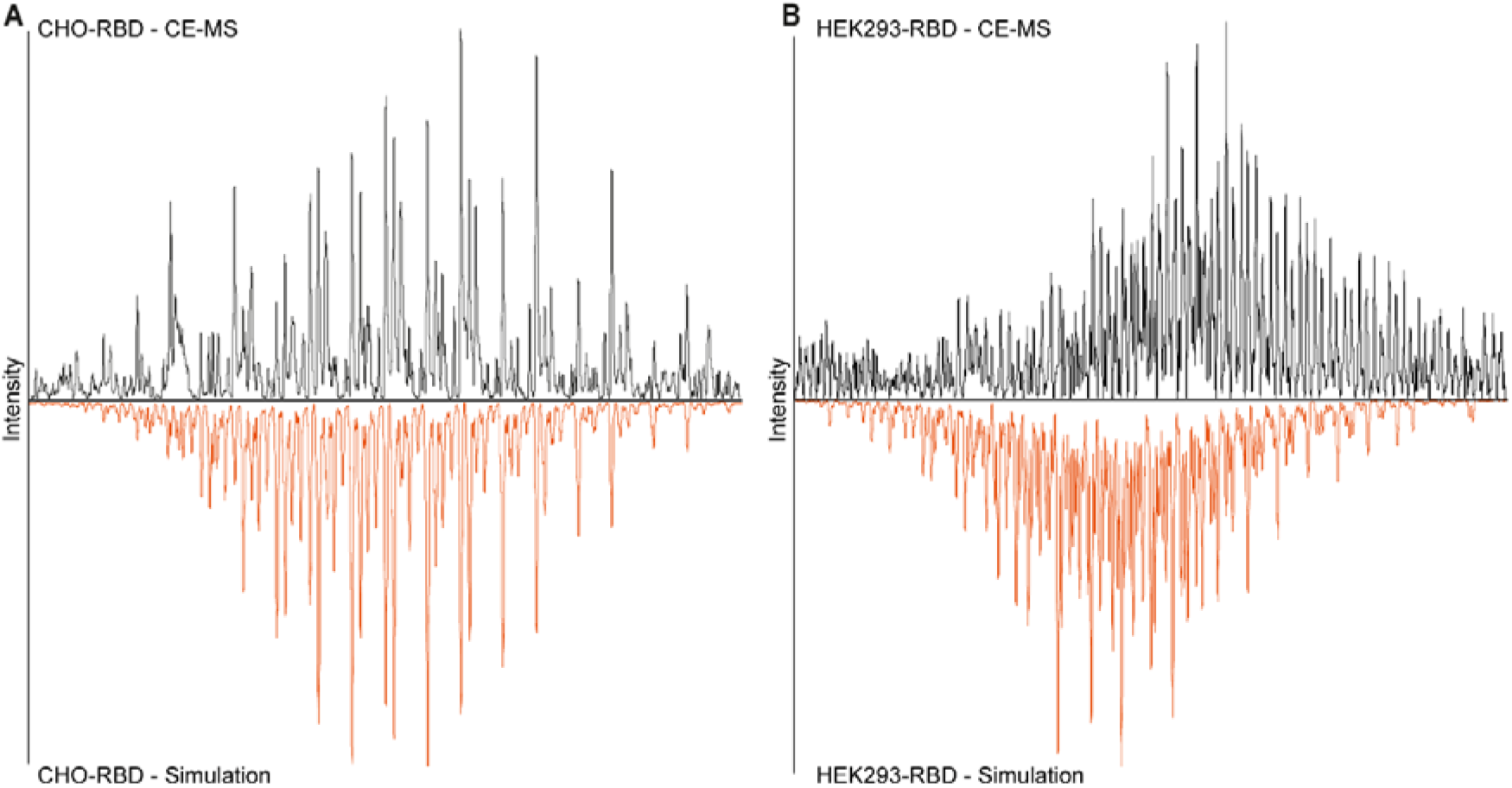
Deconvoluted mass spectra of the RBD intact (non enzymatical treated) (black trace) and *in silico* simulated mass spectrum (vermillon trace) of RBD produced in A) CHO cells or B) HEK293 cells.

### 3.2. Functional characterization of CHO and HEK293 produced RBD domains

Next to the structural characterization of the RBD also a functional characterization was performed. To assess the RBD functionality, an ACE2 receptor binding assay and a binding assay for anti-RBD antibodies from COVID-19 patient sera were performed.

#### 3.2.1. SARS-CoV-2 antibody binding assay

The binding of anti-SARS-CoV-2-IgG antibodies in sera taken from 12 Covid-19 patients more than 10 weeks after the onset of the first symptoms and RBD was determined using an ELISA assay. For the assay, the intact CHO- or HEK293-RBD, as well as deglycosylated RBD samples, were used. As a negative control, a pool of 10 sera collected in 2012 was used. As expected, higher absorption values were observed for the 12 COVID-19 patient sera compared to the negative control indicating binding of specific antibodies to the RBDs (**Figure 6**). Both intact RBD samples showed similar absorption values, while absorption values were slightly elevated when the deglycosylated RBDs were employed. Also, for the negative controls, the deglycosylated RBD showed elevated absorption values compared to the intact RBD. This is most probably the result of unspecific binding which may be increased after deglycosylation. This was also reflected in the correlation of the absorption values of the patient sera. The correlation of these values gained using intact RBDs and their deglycosylated version (CHO: R^2^ 09681, HEK293: 0.9755) was slightly lower than the correlation obtained using intact RBD produced by CHO and HEK293 cells (R^2^ 0.9894) or deglycosylated CHO-versus deglycosylated HEK293-RBD (R^2^ 0.9905) (**Figure S11**).

**Figure 6:**
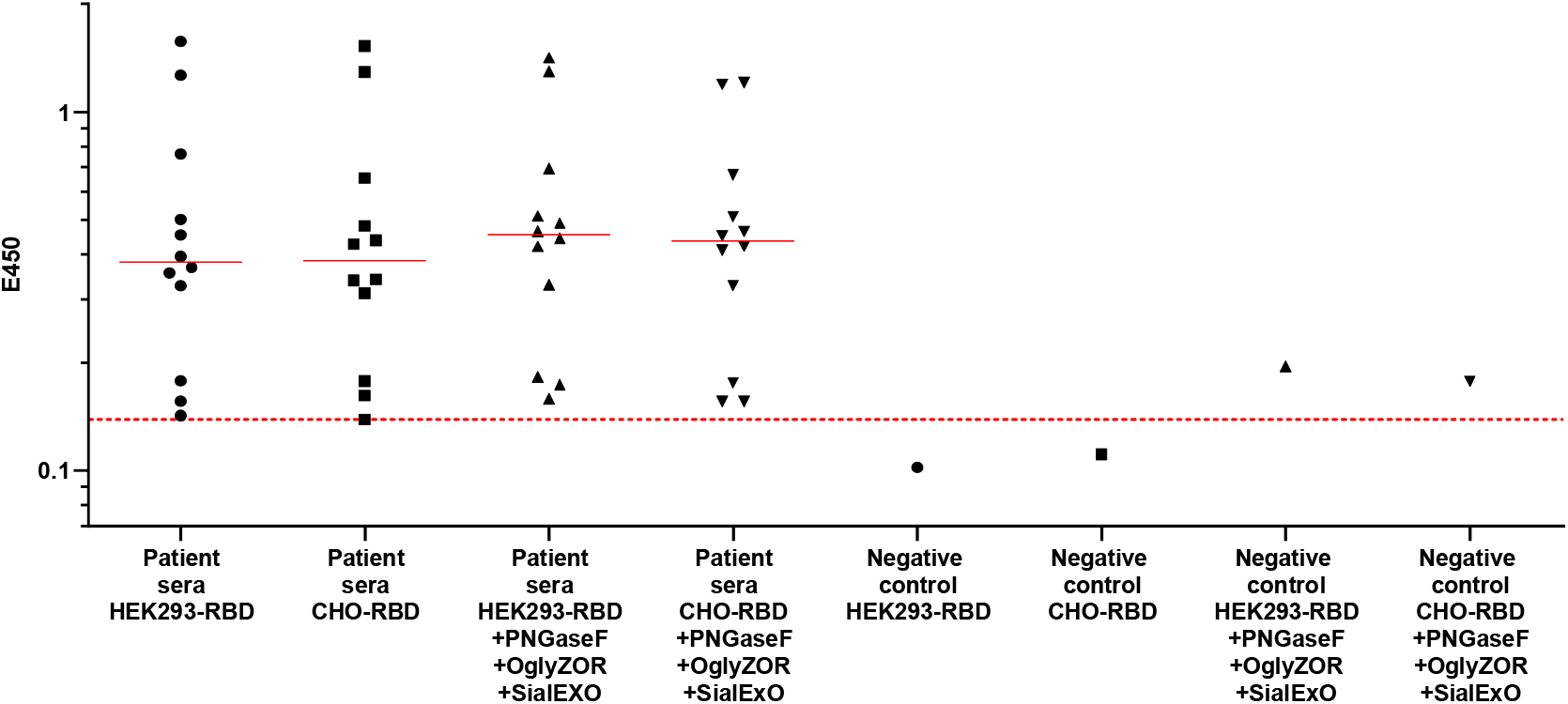
SARS-CoV-2 RBD IgG ELISA. Serum collected >10 weeks after onset of first symptoms from 12 SARS-CoV-2 PCR positive tested donors was used to test the binding to CHO and HEK293 produced intact and deglycosylated RBD samples. The negative control consists of a pool of 10 human sera from 2012. The dashed red line indicates the threshold for SARS-CoV-2 positive samples which was set to the lowest signal obtained from serum of a SARS-CoV-2 recovered patient.

#### 3.2.2 ACE2 receptor binding assay

Using a plate-based ACE2 receptor binding assay, a dose-dependent binding of both CHO- and HEK293-RBD was observed (**Figure 7A**). Comparing the RBDs expressed in both production systems, similar binding properties were observed with a trend towards a lower EC_50_ values for CHO-RBD. As glycosylation has been hypothesized to have a role in the interaction with ACE2 [10, 11], we additionally tested the RBDs after deglycosylation. Deglycosylation was found to reduce the binding of the RBDs as reflected in an approximately 2 times increase of the EC_50_ values. Of note, Qianqian *et al*. reported that the infectivity is reduced to only 0.08% after removal of the *N*-glycosylation sites [11]. Our results, although they show a slight variation between the glycosylated and non-glycosylated versions, do not explain this drastic difference in infectivity. Furthermore, the presence of endoglycosidases could also have affected the biotinylation rate resulting in the overall reduced binding. Supporting the hypothesis of Qianqian, these glycans may be crucial for stabilizing the trimeric spike protein rather than influencing the binding affinity [11].

**Figure 7:**
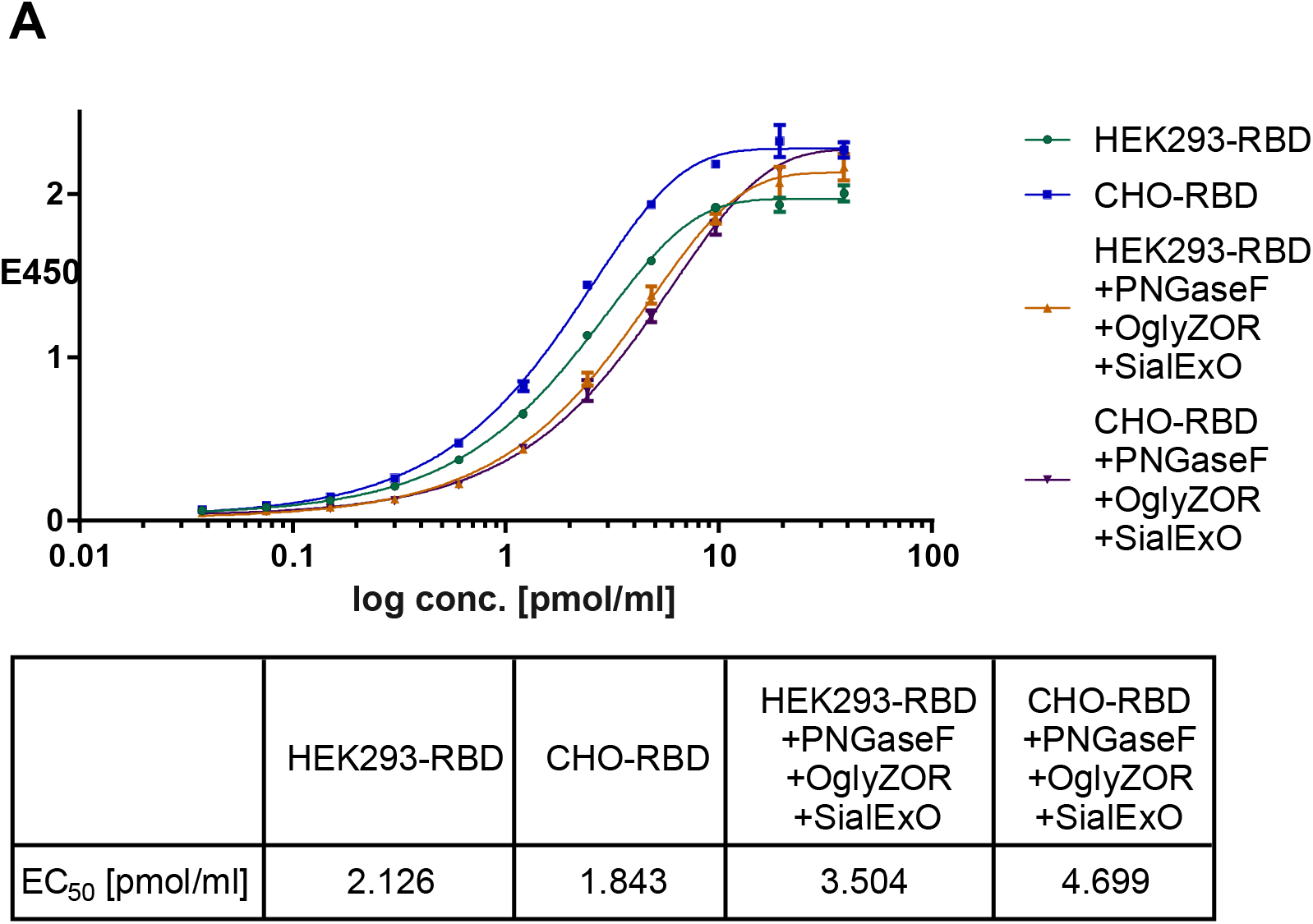
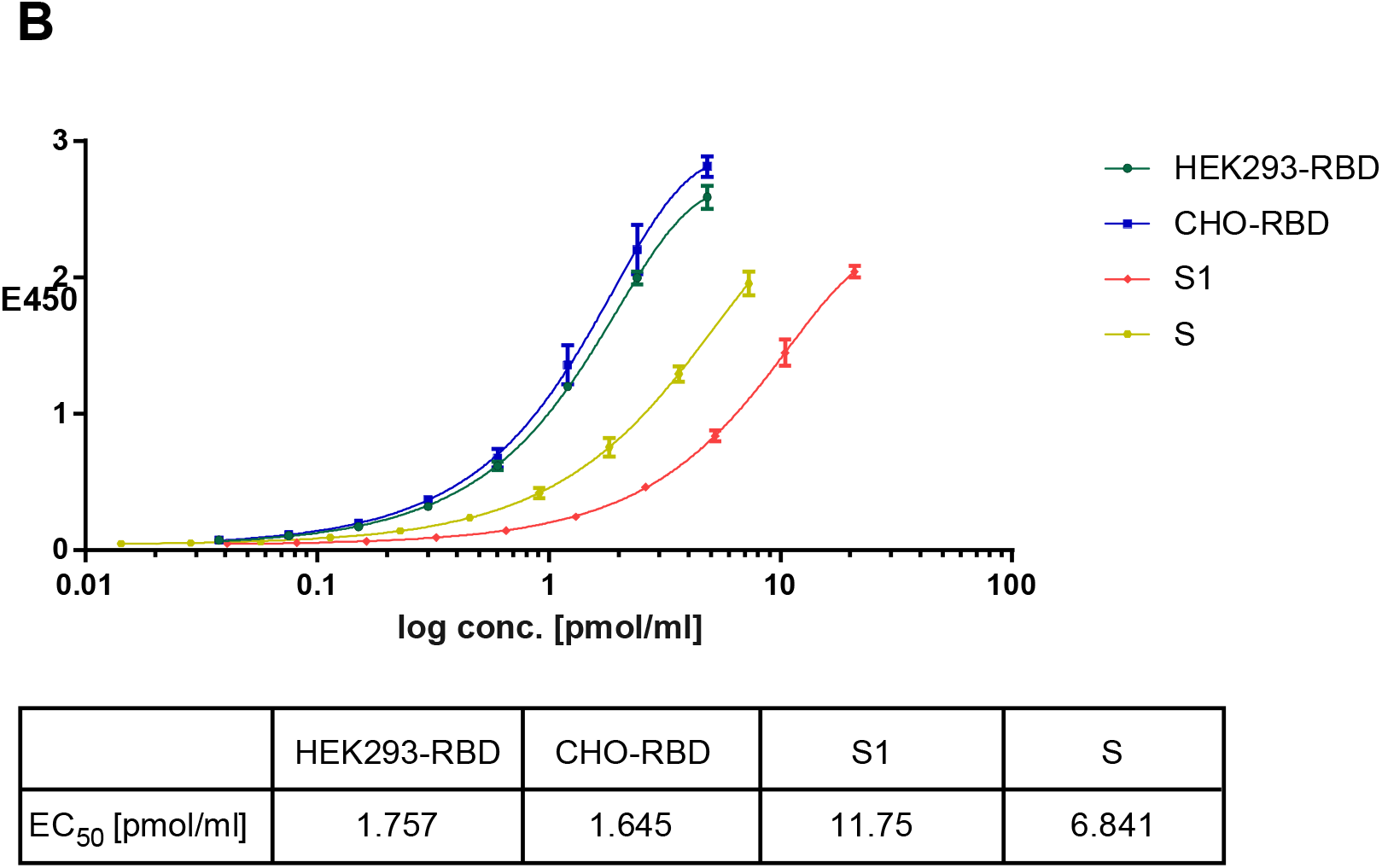
ACE2 binding assay. Dose-response curves of RBD produced in CHO or HEK293 cells binding to the ACE2 receptor. Comparison to A) deglycosylated version of CHO- or HEK293-RBD and B) S or S1 subunit (produced in HEK293 cells).

Additionally, we compared the ACE2 receptor binding affinity of both RBDs to the S1 subunit, as well as the intact S protein. The binding affinity of the ACE2 receptor to the RBDs was significantly increased compared to the S1 subunit and the S protein (**Figure 7B**). This might be due to the higher accessibility to ACE2 of the RBD only compared to the S1 subunit or even the S protein. As shown by Casalino *et al*., in the trimeric S protein the RBD can be in an up or down conformation and, therefore, more or less accessible to the ACE2 binding [8]. By using the S protein or the S1 subunit, some parts of the RBD might not be accessible or less accessible for the ACE2 receptor which may explain the reduced binding affinity. These results highlight the importance of a proper selection of recombinant proteins.

## Conclusion

RBD proteoforms were comprehensively characterized by combining intact protein, glycopeptide and released glycan analysis with enzymatic glycan dissection and topdown sequencing. The combination of multiple MS workflows was fundamental for assigning the intact RBD proteoforms. In particular, glycan dissection of the intact protein using sequentially different glycosidases has shown to be very powerful to annotate complex intact RBD spectra. The glycopeptide data, next to provide sitespecific information, was used to simulate an *in silico* intact spectrum and corroborate our assignments. This approach was applied to RBD samples from both CHO and HEK293 cells. In case of the low sialylated CHO-RBD a very strong correlation was obtained. However, for the HEK293-RBD the simulated spectrum showed a clear shift to a lower mass associated with the loss of sialic acids or an ionization bias of the glycopeptides. This stresses the importance of assessing intact proteoforms to avoid skewing of the data in any direction. The observed differences in *N*-glycosylation, with higher sialylation levels and antenna fucosylation for HEK293-RBD and low sialylation levels but high-antennary and LacNAc repeat structures for CHO-RBD are typical for the two expression systems. For the *O*-glycans, CHO-RBD showed mainly core 1 type glycans while HEK293-RBD presented a combination of core 1 and core 2 type *O*-glycans. Furthermore, using alternative approaches, such as N-terminal cleavage at the *O*-glycosylation site and MALDI-ISD we localized the *O*-glycosylation site to T323, previously unknown. Following this approach, the obtained annotated spectra could be used for batch-to-batch comparison of different RBD batches or for release testing in companies producing RBD based vaccines.

From a functional point of view, both RBDs showed similar binding to antibodies from COVID-19 patient sera as well as to the ACE2 receptor. The ACE2 binding was increased compared to S protein or S1 subunit, likely due to higher accessibility. After deglycosylation, the binding of RBDs to anti-spike antibodies remained unaffected. For ACE2 a minor decrease in the EC_50_ values was observed, which does not fully explain the infectivity decrease with aglycosylation observed in previous studies. These findings suggest that glycosylation plays a role in conformational stabilization rather than affecting binding affinity. Further studies are warranted on the role of RBD glycosylation in ACE2 binding.

## Author Contributions

C.G., T.Z., A.R., S.R. and S.P. performed the measurements and processed the data. E.D.V., M.W., T.W. and D.S. conceived the idea and designed the experiments. T.W. provided RBD samples. C.G. and E.D.V. drafted the manuscript. C.G., T.Z., A.R., S.R., S.P., D.S., T.W, M.W. and E.D.V. reviewed this manuscript equally.

## Acknowledgements

We thank Eckhard Belau and Waltraud Evers for preparing the samples and analysis. This work was supported by the Analytics for Biologics project (Grant agreement ID 765502) of the European Commission.

