## Supplementary information for "Structural and functional characterization of SARS-CoV-2 RBD domains produced in mammalian cells"

*Dr. Elena Domínguez-Vega

Leiden University Medical Center, Center for Proteomics and Metabolomics, Albinusdreef 2. 2333ZA Leiden, The Netherlands.

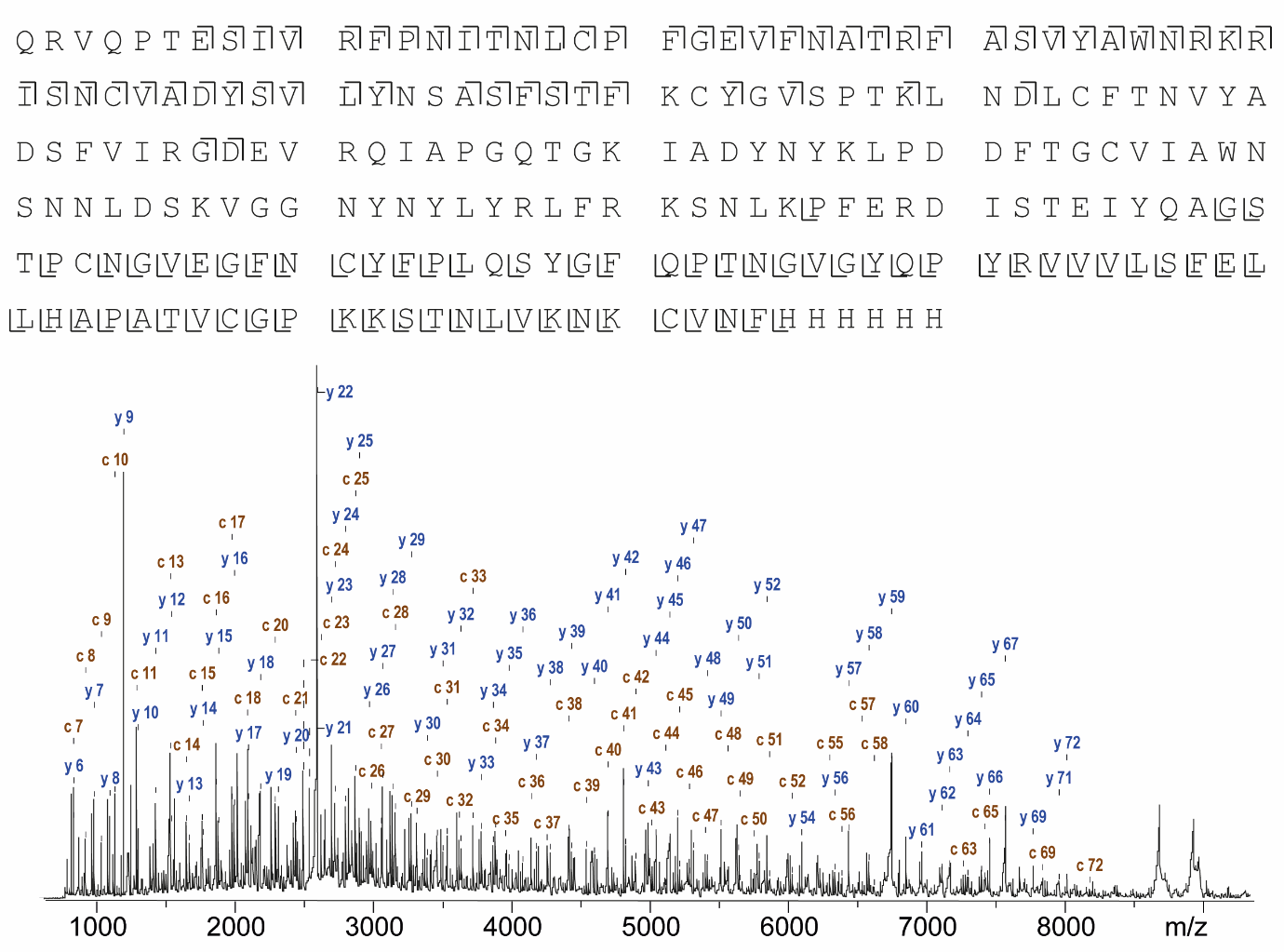

**Figure S1:** Sequence coverage observed with MALDI-ISD Top-down of HEK293-RBD after PNGAseF and OglyZOR treatment and reduction. For assignemt a and c ions in case of the N-terminus or y and z+2 ions for the C-terminus were used. In the MALDI-ISD spectrum N-terminal c-ions are shown in brown and the C-terminal y-ions are shown in blue.

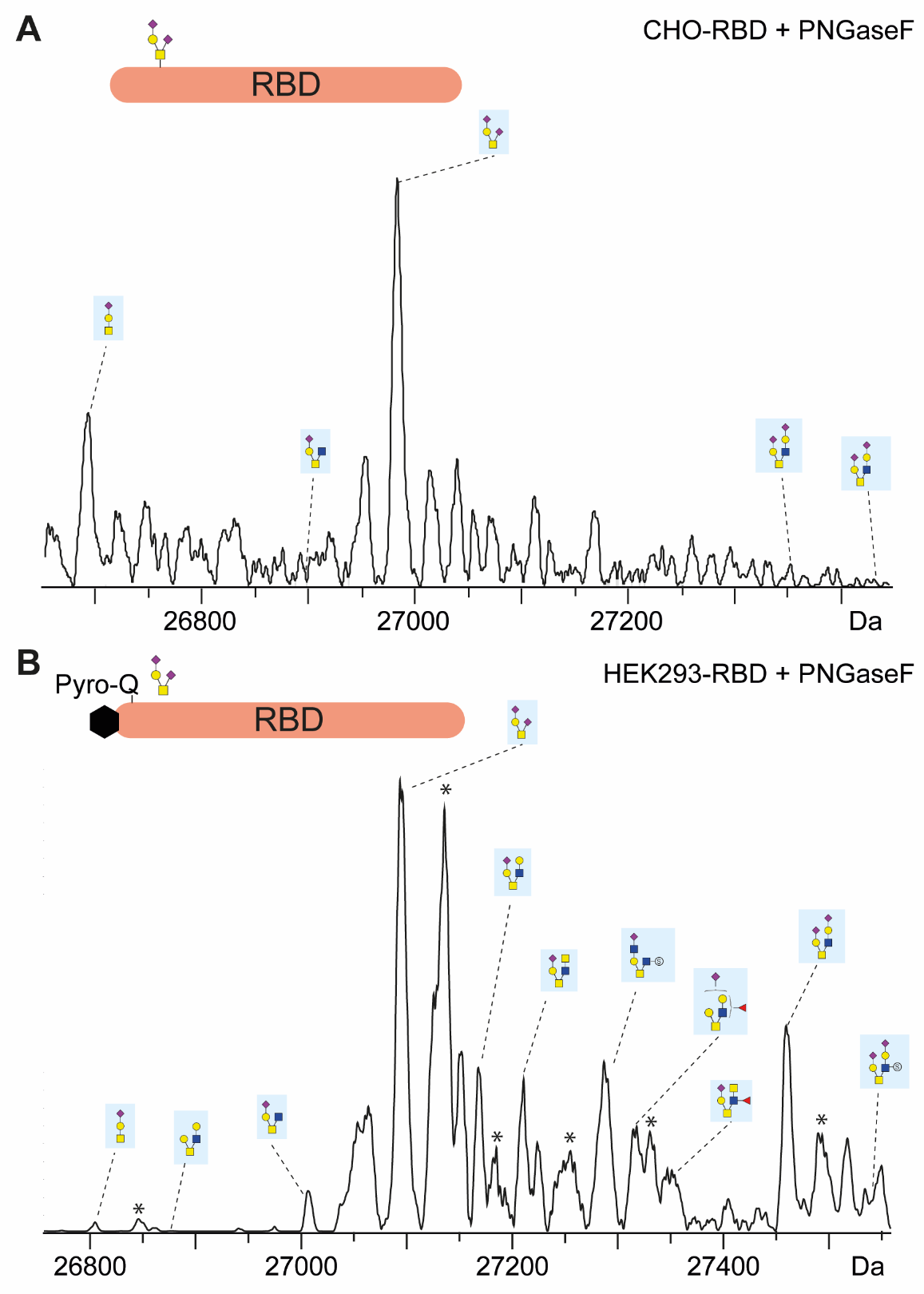

**Figure S2**: Deconvoluted ESI mass spectra of the RBD treated with PNGaseF produced by A) CHO cells or B) HEK293 cells. Peaks marked with an asterisk * presumably correspond to the acetylated variant of RBD. Yellow square, N-acetylgalactosamine; yellow circle, galactose; blue square, N-acetylglucosamine; red triangle, fucose; purple diamond, N-acetylneuraminic acid (sialic acid), S, sulfate.

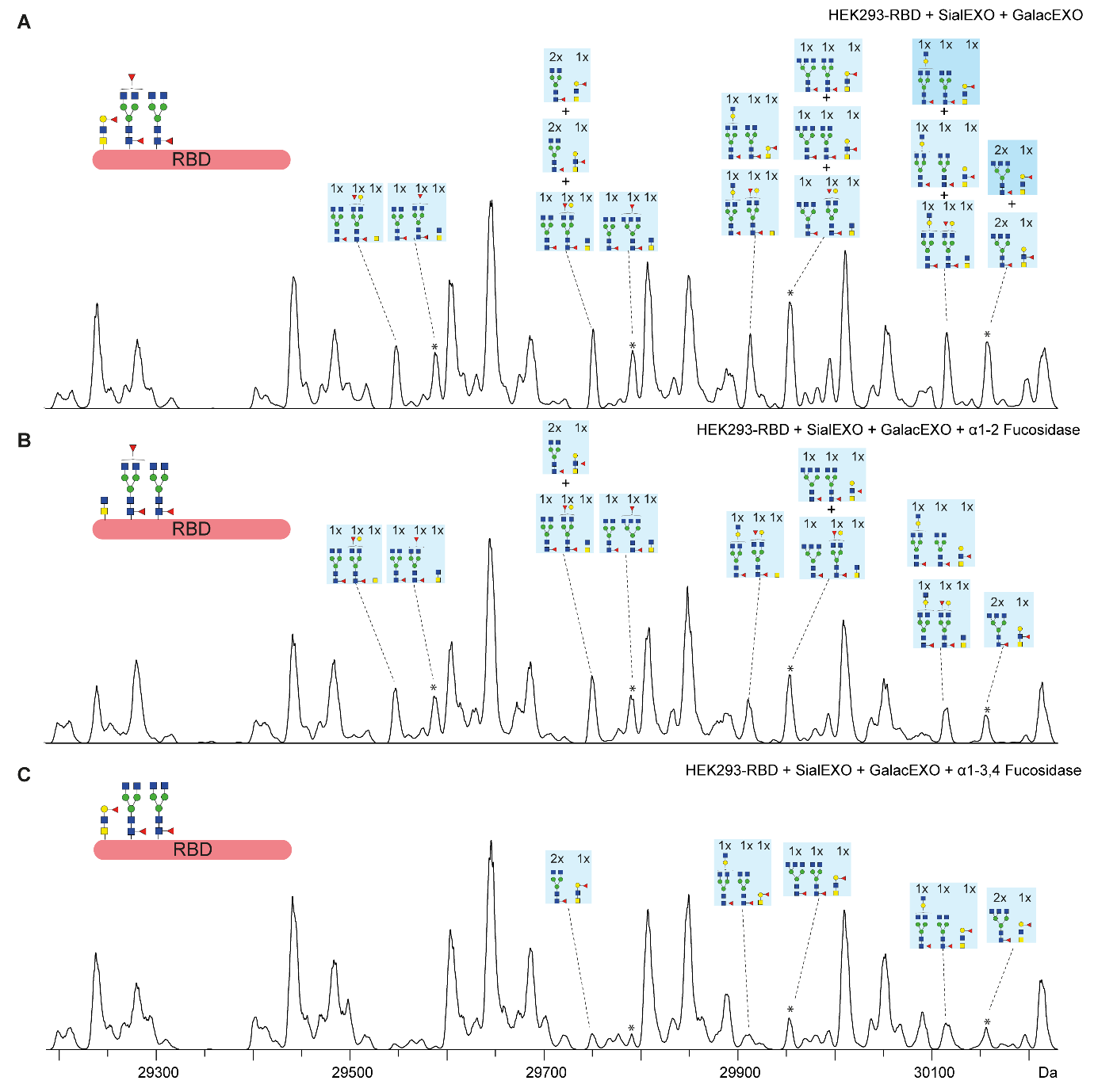

**Figure S3**: Deconvoluted ESI mass spectra of the HEK293-RBD treated with either A) SialEXO and GalactEXO, B) SialEXO, GalactEXO and α1-2 fucosidase or C) SialEXO, GalactEXO and α1-3,4 fucosidase. For simplicity only antenna of fucosylated glycans are assigned. For complete assignment compare figure S5B. Peaks marked with an asterisk * contain additionally the acetylated variant of RBD. Yellow square, N-acetylgalactosamine; yellow circle, galactose; blue square, N-acetylglucosamine; red triangle, fucose.

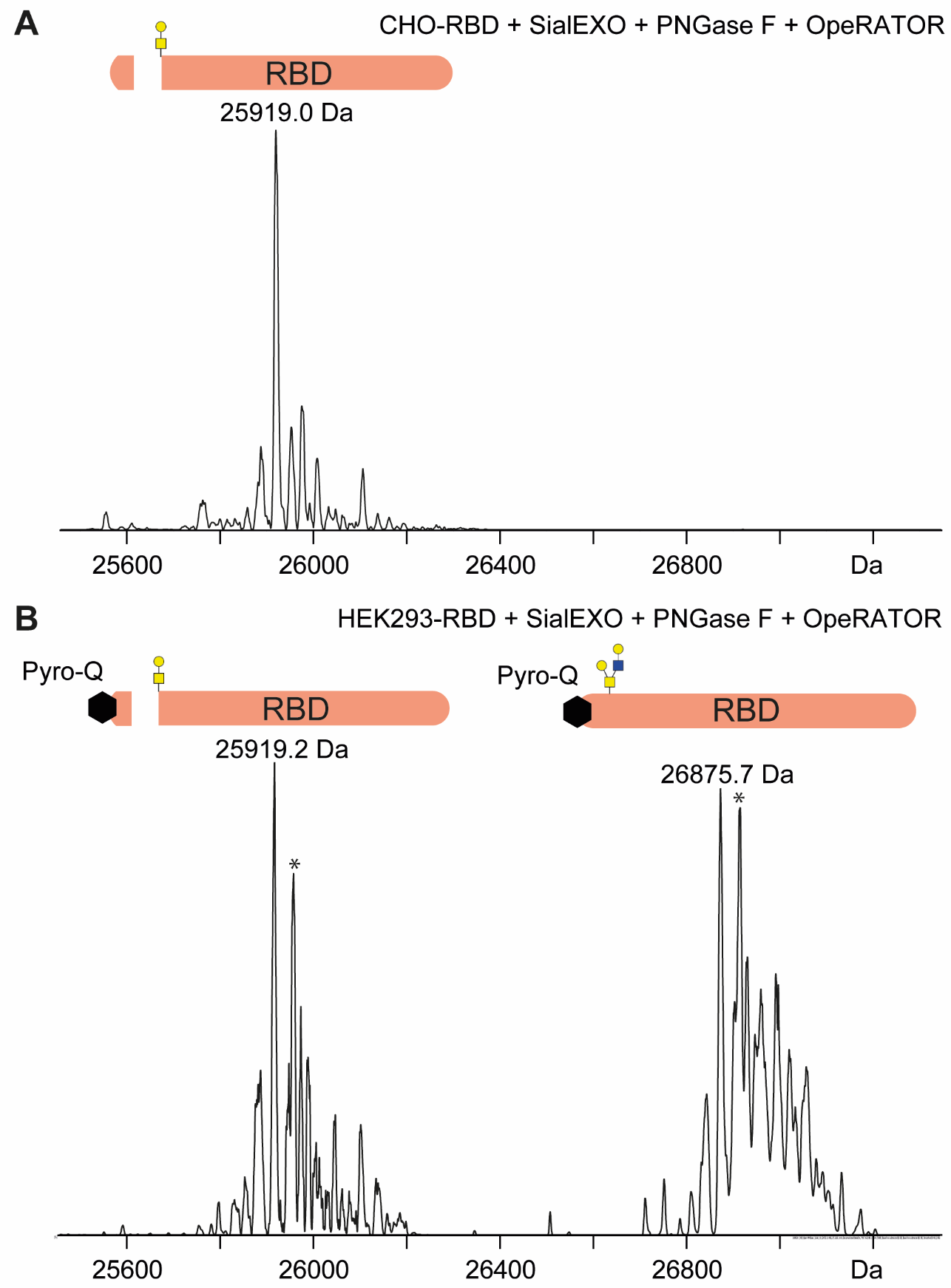

**Figure S4**: Deconvoluted ESI mass spectra of the RBD produced by A) HEK293 cells or B) CHO cells treated with PNGaseF, SialEXO and OpeRATOR. Peaks marked with an asterisk * presumably correspond to the acetylated variant of RBD. Yellow square, N-acetylgalactosamine; yellow circle, galactose; blue square, N-acetylglucosamine.

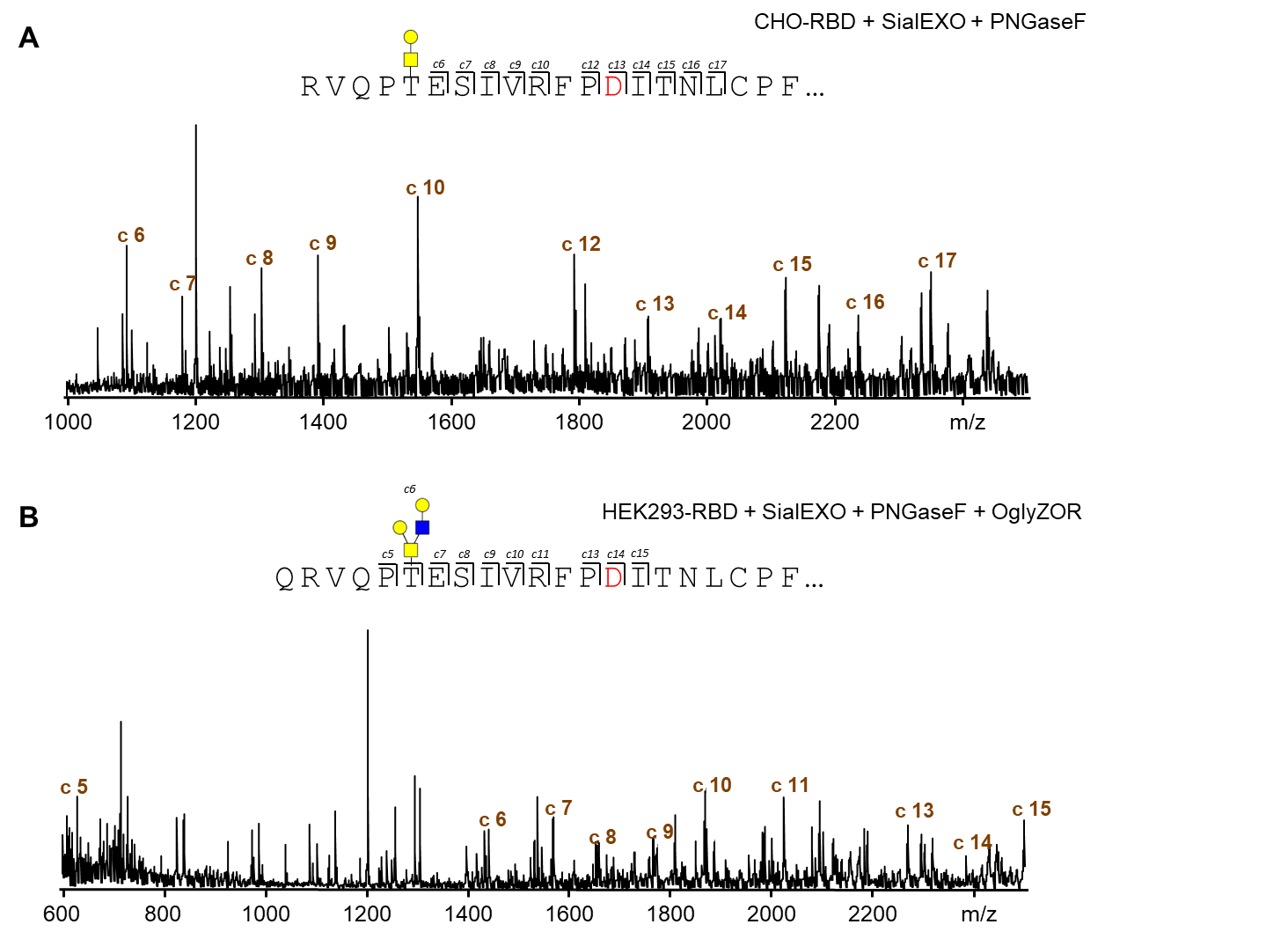

**Figure S5**: MALDI-ISD mass spectra. A) Detected c-ions of CHO-RBD after SialEXO and PNGaseF treatment. B) Detected c-ions of HEK293-RBD after SialEXO, PNGaseF and OglyZOR treatment. T* indicates a matching O-glycosylation at threonine. Yellow square, N-acetylgalactosamine; yellow circle, galactose; blue square, N-acetylglucosamine.

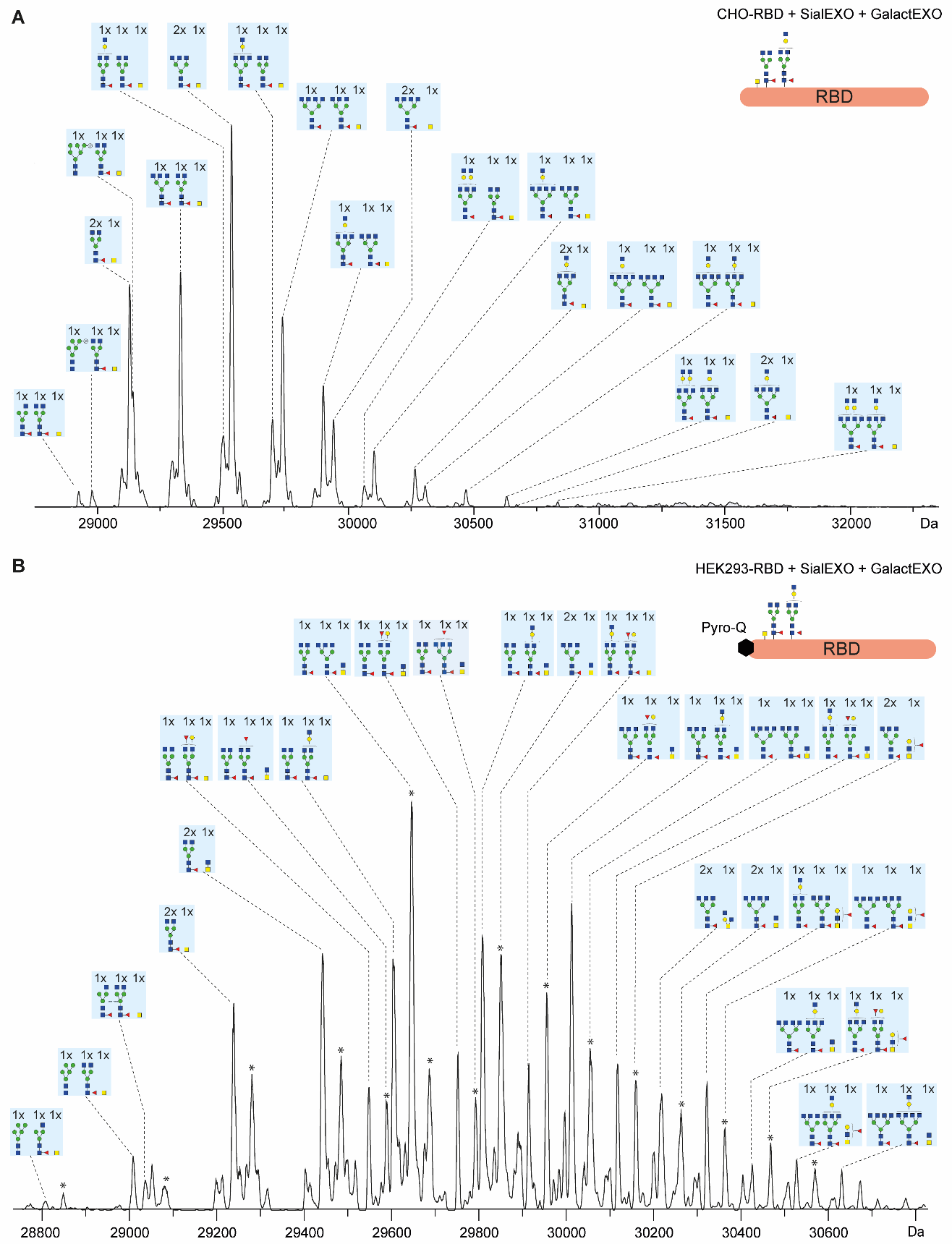

**Figure S6**: Deconvoluted ESI mass spectra of the CHO-RBD treated with A) SialEXO and SialEXO + GalactEXO. The assignments were based on observed mass, enzymatic treatment and in case of identical mass on the most likely combination using the intensities obtained by the glycopeptide data. Yellow square, N-acetylgalactosamine; yellow circle, galactose; blue square, N-acetylglucosamine; red triangle, fucose.

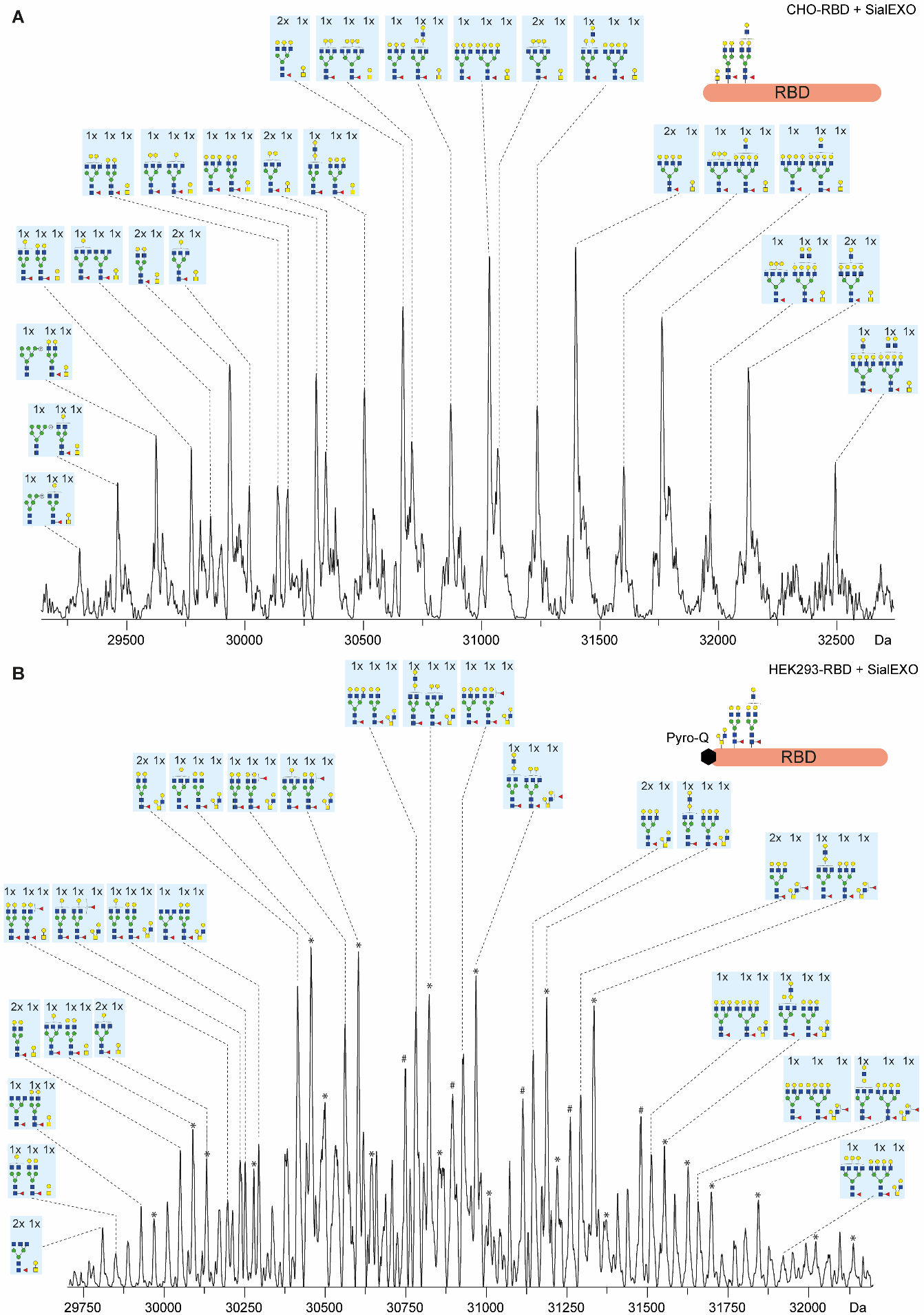

**Figure S7**: Deconvoluted ESI mass spectra of the HEK293-RBD treated with A) SialEXO and SialEXO + GalactEXO. The assignments were based on observed mass, enzymatic treatment and in case of identical mass on the most likely combination using the intensities obtained by the glycopeptide data. Peaks marked with an asterisk * are due to the acetylated variant of RBD. Yellow square, N-acetylgalactosamine; yellow circle, galactose; blue square, N-acetylglucosamine; red triangle, fucose.

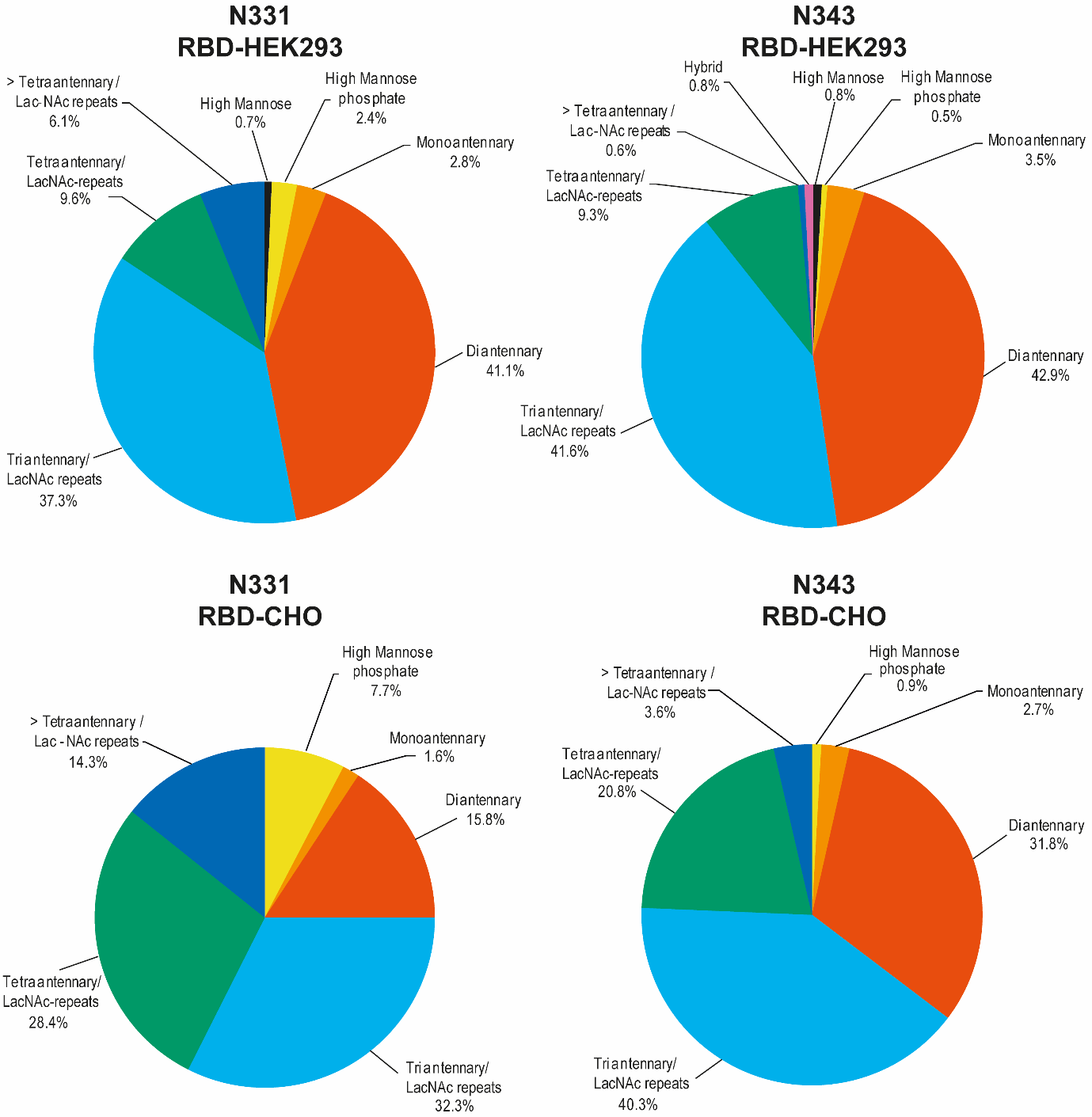

**Figure S8**: Pie charts visualizing the glycan type distribution for the different glycosylation sites (N331 and N343) and the different RBD samples (HEK293 or CHO). Data are based on Tables S2 and S3.

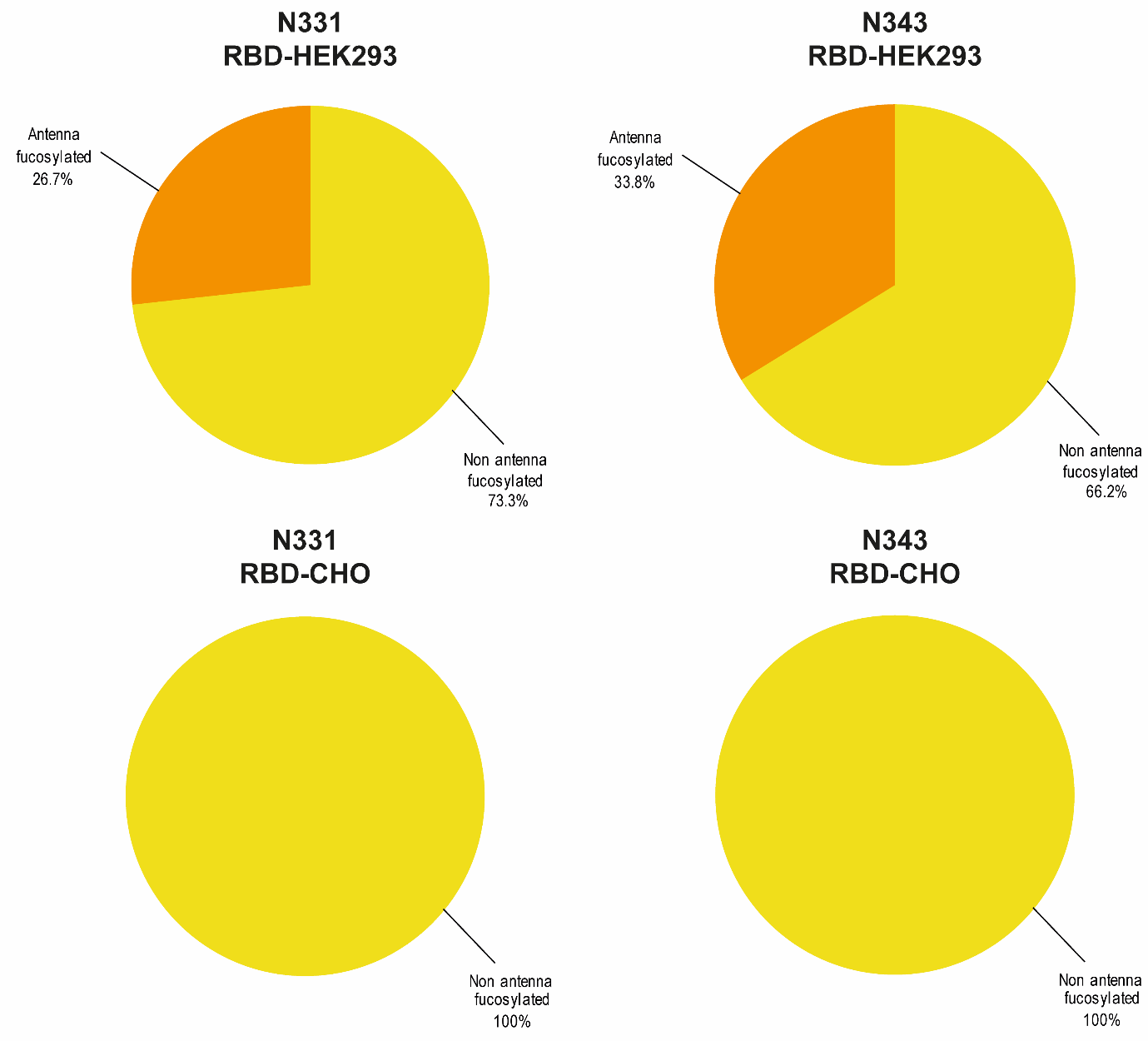

**Figure S9**: Pie charts visualizing the amount of antenna fucosylation for the different glycosylation sites (N331 and N343) and the different RBD samples (HEK293 or CHO). Data are based on Tables S2 and S3.

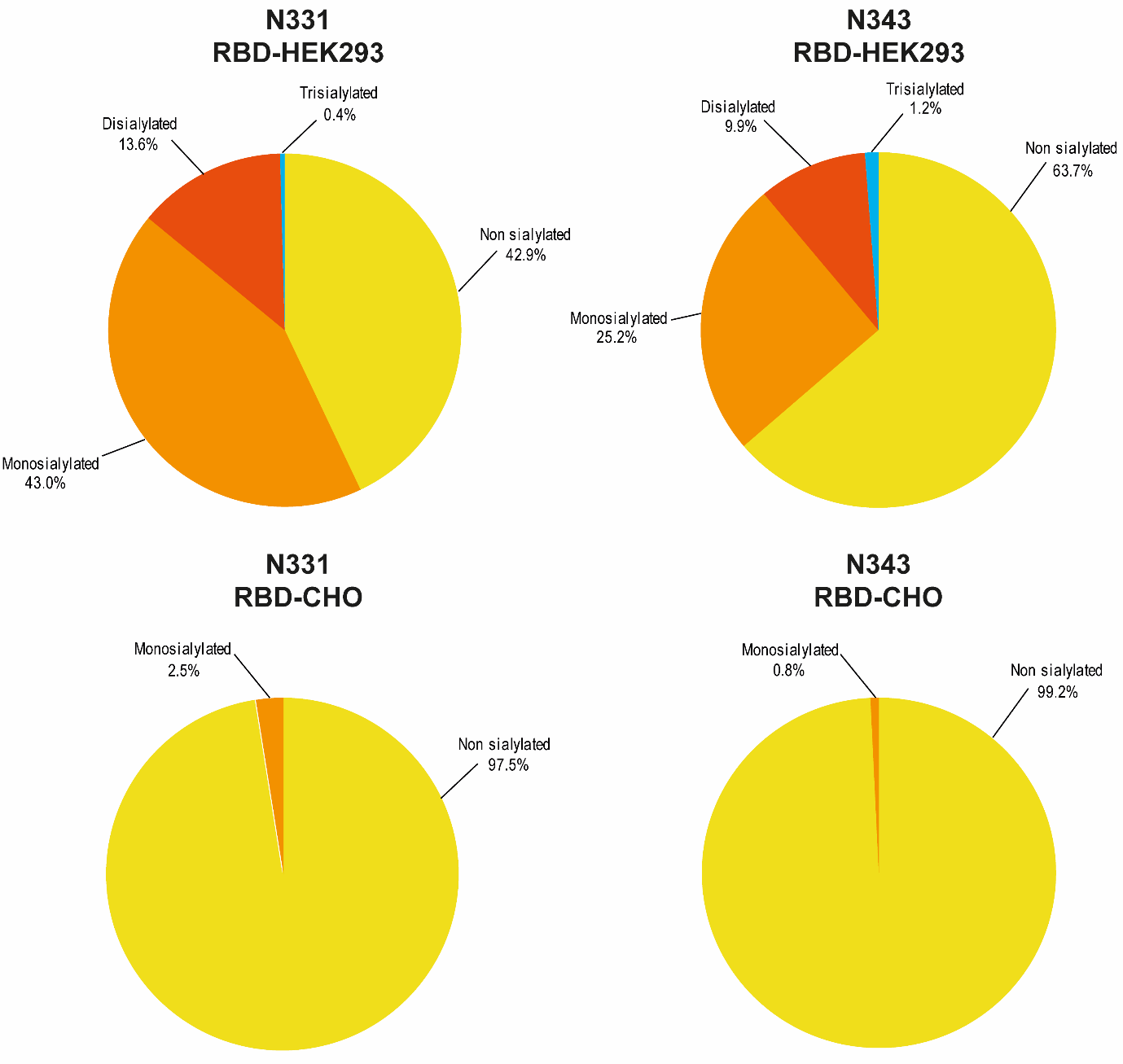

**Figure S10**: Pie charts visualizing the level of sialylation for the different glycosylation sites (N331 and N343) for samples HEK293- and CHO-RBD. Data are based on Tables S2 and S3.

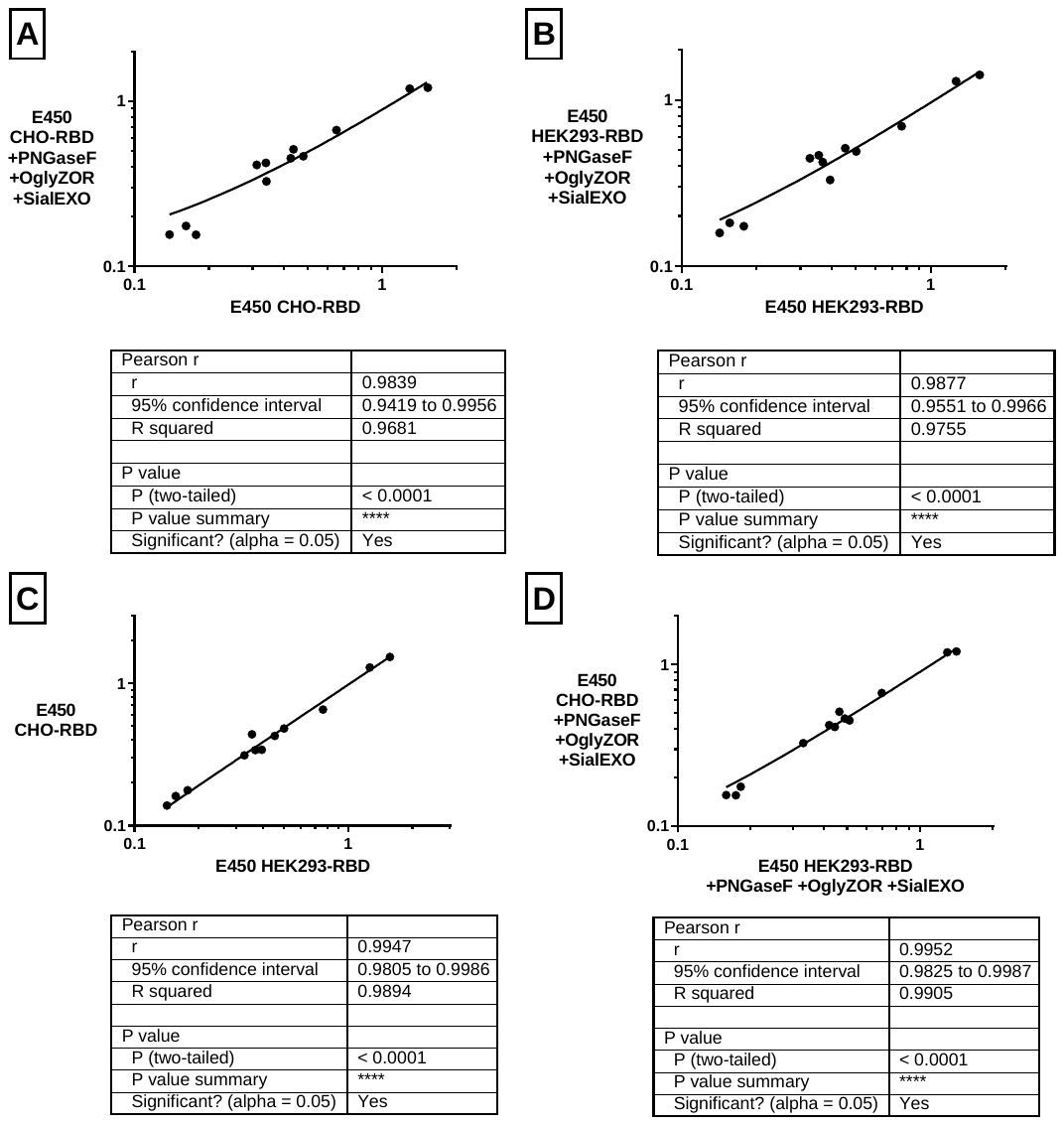

**Figure S11**: Correlation analysis of absorption values of 12 different sera from SARS-CoV-2-IgG positive patients obtained using an ELISA with differently glycosylated RBDs as coating antigens. A) CHO-RBD and deglycosylated CHO-RBD, B) HEK293-RBD and deglycosylated HEK293-RBD, C) CHO-RBD and HEK293-RBD and D) deglycosylated CHO- and deglycosylated HEK293-RBD.

**Table S1**. Relative quantification of *O*-glycans released from RBD using PGC nano-LC-ESI-MS/MS. The composition of *O*-glycans have been assigned based on mass and fragmentation spectra. H: hexose; N: *N*-acetylhexosamine; F: fucose; S: *N*-acetylneuraminic acid; Su: sulphate. n.d.: not determined.

| **Composition** | **Rel. abundance (%)**  **CHO HEK** | | **Observed ions** | | | **Theoretical** | **Deviation** |
| --- | --- | --- | --- | --- | --- | --- | --- |
|  |  |  | **[M-H]^-^** | **[M-2H]^2-^** | | **[M-H]^-^** | **Δ[M-H]^-^** |
| H1N1 | 1.18 | 0 | 384.13 | n.d. |  | 384.151 | 0.021 |
| N1S1 | 0.56 | 0.07 | 513.13 | n.d. |  | 513.194 | 0.064 |
| H1N2 | 0 | 0.15 | 587.17 | n.d. |  | 587.231 | 0.061 |
| H1N1S1 | 30.03 | 2.32 | 675.21 | n.d. |  | 675.247 | 0.037 |
| H2N2 | 0.12 | 0.63 | 749.24 | n.d. |  | 749.283 | 0.043 |
| H1N3 | 0 | 0.39 | 790.27 | n.d. |  | 790.310 | 0.040 |
| H1N1S1F1 | 0 | 0.30 | 821.26 | n.d. |  | 821.304 | 0.044 |
| H2N2Su | 0.68 | 0.35 | 829.25 | n.d. |  | 829.240 | 0.010 |
| H1N2S1 | 0.79 | 7.08 | 878.28 | n.d. |  | 878.326 | 0.046 |
| H2N2F1 | 0 | 0.19 | 895.29 | n.d. |  | 895.341 | 0.051 |
| H1N1S2 | 59.24 | 26.63 | 966.33 | n.d. |  | 966.342 | 0.012 |
| H2N2S1 | 1.02 | 9.90 | 1040.36 | n.d. |  | 1040.379 | 0.019 |
| H1N3S1 | 0.35 | 3.31 | 1081.37 | n.d. |  | 1081.405 | 0.035 |
| H1N3S1Su | 1.56 | 14.46 | 1161.32 | n.d. |  | 1161.362 | 0.042 |
| H2N2S1F1 | 0.41 | 4.67 | 1186.40 | n.d. |  | 1186.437 | 0.037 |
| H1N3S1F1 | 0.10 | 1.37 | 1227.42 | n.d. |  | 1227.463 | 0.043 |
| H2N2S1F1Su | 0 | 0.46 | 1266.34 | n.d. |  | 1266.394 | 0.054 |
| H2N2S2 | 3.09 | 20.27 | 1331.40 | 665.20 |  | 1331.474 | 0.034 |
| H2N2S1F2 | 0.10 | 0.87 | 1332.44 | n.d. |  | 1332.495 | 0.055 |
| H2N2S2Su | 0.76 | 6.58 | 1411.39 | 705.15 |  | 1411.431 | 0.041 |

**Table S2.** Glycopeptides for the *N*-glycosylation site (N343) from RBD using a trypsin and elastase digestion and separation by LC-ESI-MS/MS. The composition of *N*-glycans have been assigned based on mass and fragmentation spectra. H: hexose; N: *N*-acetylhexosamines; F: fucose; S: *N*-acetylneuraminic acid; G: *N*-Glycolylneuraminic acid; P: phosphate; n.d.: not determined. Aglycon: GEVFNATR.

| **Composition** | | **Rel. abundance (%)** | | | **Observed ions** | | **Theoretical** | | | **Deviation** | |
| --- | --- | --- | --- | --- | --- | --- | --- | --- | --- | --- | --- |
|  | **HEK293** | | **CHO** | **[M+H]^+^** | | **[M+2H]^2+^** | | **[M+3H]^3+^** | **[M+2H]^2+^** | | **Δ[M+2H]^2+^** |
| H3N2F1 | 0.08 | | 0.13 | n.d. | | 966.922 | | n.d. | 966.918 | | 0.004 |
| H6N2 | 0.55 | | 0 | n.d. | | 1136.970 | | n.d. | 1136.971 | | 0.001 |
| H7N2 | 0.21 | | 0 | n.d. | | 1217.997 | | n.d. | 1217.995 | | 0.002 |
| H8N2 | 0.07 | | 0 | n.d. | | 1299.023 | | n.d. | 1299.022 | | 0.001 |
| H5N2P1 | 0 | | 0.05 | n.d. | | 1095.934 | | n.d. | 1095.932 | | 0.002 |
| H6N2P1 | 0.34 | | 0.76 | n.d. | | 1176.956 | | n.d. | 1176.958 | | 0.002 |
| H7N2P1 | 0.20 | | 0.07 | n.d. | | 1257.982 | | n.d. | 1257.985 | | 0.001 |
| H3N3F1 | 1.16 | | 1.50 | n.d. | | 1068.460 | | n.d. | 1068.460 | | 0.000 |
| H4N3F1 | 0.71 | | 1.03 | n.d. | | 1149.487 | | n.d. | 1149.487 | | 0.000 |
| H4N3F2 | 0.97 | | 0 | n.d. | | 1222.515 | | 815.347 | 1222.516 | | 0.000 |
| H4N3S1F1 | 0.56 | | 0 | n.d. | | 1295.033 | | n.d. | 1295.035 | | 0.002 |
| H5N3F1 | 0.56 | | 0 | n.d. | | 1230.512 | | n.d. | 1230.513 | | 0.002 |
| H6N3F1 | 0.23 | | 0 | n.d. | | 1311.037 | | n.d. | 1311.033 | | 0.002 |
| H3N4F1 | 5.05 | | 5.21 | n.d. | | 1169.999 | | 780.337 | 1170.000 | | 0.002 |
| H4N4F1 | 3.06 | | 10.16 | n.d. | | 1251.026 | | 834.355 | 1251.026 | | 0.000 |
| H4N4F2 | 2.79 | | 0 | n.d. | | 1324.057 | | 883.039 | 1324.058 | | 0.002 |
| H4N4S1F1 | 2.43 | | 0 | n.d. | | 1396.573 | | 931.387 | 1396.576 | | 0.003 |
| H5N4F1 | 0 | | 15.93 | n.d. | | 1332.053 | | 888.372 | 1332.054 | | 0.001 |
| H5N4F2 | 5.78 | | 0 | n.d. | | 1405.084 | | 937.056 | 1405.083 | | 0.001 |
| H5N4F3 | 3.81 | | 0 | n.d. | | 1478.111 | | 985.745 | 1478.111 | | 0.000 |
| H5N4S1F1 | 7.14 | | 0.29 | n.d. | | 1477.603 | | 985.403 | 1477.600 | | 0.004 |
| H5N4G1F1 | 0.29 | | 0.00 | n.d. | | 1485.586 | | n.d. | 1485.594 | | 0.009 |
| H5N4S1F2 | 6.47 | | 0 | n.d. | | 1550.632 | | 1034.089 | 1550.629 | | 0.004 |
| H5N4S2F1 | 5.78 | | 0 | n.d. | | 1623.151 | | 1082.435 | 1623.148 | | 0.003 |
| H6N4F1 | 0.17 | | 0.20 | n.d. | | 1413.081 | | 942.388 | 1413.079 | | 0.003 |
| H6N4F2 | 0.09 | | 0 | n.d. | | 1486.108 | | n.d. | 1486.105 | | 0.003 |
| H3N5F1 | 8.48 | | 5.88 | n.d. | | 1271.541 | | 848.028 | 1271.539 | | 0.001 |
| H4N5F1 | 6.21 | | 11.08 | n.d. | | 1352.568 | | 902.048 | 1352.568 | | 0.000 |
| H4N5F2 | 4.23 | | 0 | n.d. | | 1425.594 | | 950.734 | 1425.595 | | 0.001 |
| H4N5S1F1 | 3.25 | | 0.24 | n.d. | | 1498.113 | | 998.746 | 1498.114 | | 0.001 |
| H5N5S2F2 | 2.25 | | 0 | n.d. | | 1571.142 | | 1047.766 | 1571.142 | | 0.000 |
| H5N5F1 | 3.15 | | 12.16 | n.d. | | 1433.592 | | 956.066 | 1433.593 | | 0.002 |
| H5N5F2 | 1.62 | | 0 | n.d. | | 1506.621 | | 1004.752 | 1506.621 | | 0.000 |
| H5N5S1F1 | 1.85 | | 0 | n.d. | | 1579.143 | | 1053.096 | 1579.141 | | 0.002 |
| H6N5F1 | 2.32 | | 10.74 | n.d. | | 1514.621 | | 1010.082 | 1514.619 | | 0.002 |
| H6N5F2 | 1.79 | | 0 | n.d. | | 1587.648 | | 1058.770 | 1587.650 | | 0.002 |
| H6N5F3 | 0.90 | | 0 | n.d. | | 1660.679 | | 1107.454 | 1660.677 | | 0.002 |
| H6N5S1F1 | 2.47 | | 0.22 | n.d. | | 1660.170 | | 1107.114 | 1660.167 | | 0.003 |
| H6N5S2F1 | 1.88 | | 0 | n.d. | | 1805.717 | | 1204.146 | 1805.714 | | 0.002 |
| H6N5S3F1 | 1.21 | | 0 | n.d. | | 1950.766 | | 1300.844 | 1950.762 | | 0.004 |
| H3N6 | 1.50 | | 0 | n.d. | | 1299.560 | | 866.712 | 1299.556 | | 0.003 |
| H3N6F1 | 1.94 | | 0.97 | n.d. | | 1373.081 | | 915.722 | 1373.079 | | 0.003 |
| H3N6F2 | 1.52 | | 0 | n.d. | | 1446.108 | | 964.410 | 1446.106 | | 0.002 |
| H3N6S1F1 | 0.50 | | 0 | n.d. | | 1518.634 | | n.d. | 1518.6283 | | 0.006 |
| H3N6S1F2 | 0.23 | | 0 | n.d. | | 1591.658 | | 1061.440 | 1591.654 | | 0.004 |
| H4N6F1 | 0.99 | | 2.42 | n.d. | | 1454.108 | | 969.739 | 1454.102 | | 0.006 |
| H4N6F2 | 0.57 | | 0 | n.d. | | 1527.137 | | 1018.423 | 1527.137 | | 0.000 |
| H5N6F1 | 1.01 | | 3.84 | n.d. | | 1535.135 | | 1023.757 | 1535.128 | | 0.006 |
| H5N6F2 | 0.68 | | 0 | n.d. | | 1608.160 | | 1072.444 | 1608.161 | | 0.001 |
| H6N6F1 | 0.12 | | 5.94 | n.d. | | 1616.161 | | 1077.775 | 1616.155 | | 0.005 |
| H6N6F2 | 0.10 | | 0.00 | n.d. | | 1688.691 | | 1126.127 | 1688.687 | | 0.004 |
| H7N6F1 | 0.11 | | 7.56 | n.d. | | 1697.190 | | 1131.792 | 1697.183 | | 0.007 |
| H3N7F1 | 0.32 | | 0 | n.d. | | 1474.621 | | 983.415 | 1474.618 | | 0.003 |
| H4N7F1 | 0.20 | | 0 | n.d. | | 1555.649 | | 1037.432 | 1555.645 | | 0.004 |
| H6N7F1 | 0 | | 0.56 | n.d. | | 1717.701 | | 1145.469 | 1717.701 | | 0.000 |
| H7N7F1 | 0 | | 0.93 | n.d. | | 1798.225 | | 1199.152 | 1798.226 | | 0.001 |
| H8N7F1 | 0 | | 2.13 | n.d. | | 1879.753 | | 1253.504 | 1879.751 | | 0.002 |
| H3N8F1 | 0.10 | | 0 | n.d. | | 1576.162 | | 1051.108 | 1576.159 | | 0.003 |

**Table S3.** Glycopeptides for the *N*-glycosylation site (N331) from RBD using a trypsin and elastase digestion and separation by LC-ESI-MS/MS. The composition of *N*-glycans have been assigned based on mass and fragmentation spectra. H: hexose; N: *N*-acetylhexosamines; F: fucose; S: *N*-acetylneuraminic acid; P: phosphate; n.d.: not determined. Aglycon: FPNIT.

| **Composition** | **Rel. abundance (%)** | | | | **Observed ions** | | | **Theoretical** | **Deviation** |
| --- | --- | --- | --- | --- | --- | --- | --- | --- | --- |
|  | | **HEK293** | | **CHO** | **[M+H]+** | **[M+2H]^2+^** | **[M+3H]^3+^** | **[M+2H]^2+^** | **Δ[M+2H]^2+^** |
| H5N2 | | | 0.56 | 0 | n.d. | 904.377 | n.d. | 904.372 | 0.005 |
| H6N2 | | | 0.07 | 0 | n.d. | 985.404 | n.d. | 985.402 | 0.003 |
| H7N2 | | | 0.07 | 0 | n.d. | 1066.425 | n.d. | 1066.431 | 0.006 |
| H5N2P1 | | | 0.40 | 2.59 | 1887.711 | 944.360 | n.d. | 944.358 | 0.002 |
| H6N2P1 | | | 1.79 | 5.09 | 2050.768 | 1025.388 | n.d. | 1025.388 | 0.000 |
| H7N2P1 | | | 0.19 | 0 | n.d. | 1106.413 | n.d. | 1106.414 | 0.001 |
| H3N3F1 | | | 1.81 | 0 | 1832.776 | 916.892 | n.d. | 916.889 | 0.003 |
| H4N3F1 | | | 0.37 | 1.62 | 1994.827 | 997.919 | n.d. | 997.914 | 0.005 |
| H4N3S1F1 | | | 0.62 | 0 | n.d. | 1143.969 | n.d. | 1143.969 | 0.001 |
| H3N4F1 | | | 3.86 | 0.29 | 2036.860 | 1018.432 | n.d. | 1018.428 | 0.004 |
| H3N4F2 | | | 0.93 | 0 | 2182.921 | 1091.466 | n.d. | 1091.457 | 0.009 |
| H4N4F1 | | | 0 | 5.19 | 2197.910 | 1099.459 | n.d. | 1099.454 | 0.005 |
| H4N4F2 | | | 1.98 | 0 | 2344.971 | 1172.990 | n.d. | 1172.987 | 0.003 |
| H4N4S1F1 | | | 4.45 | 0 | n.d. | 1245.509 | 830.674 | 1245.507 | 0.002 |
| H5N4F1 | | | 0 | 10.29 | n.d. | 1180.985 | 787.661 | 1180.988 | 0.003 |
| H5N4F2 | | | 2.49 | 0 | 2507.026 | 1254.017 | n.d. | 1254.014 | 0.002 |
| H5N4S1F1 | | | 10.69 | 0 | n.d. | 1326.535 | 884.693 | 1326.533 | 0.001 |
| H5N4S1F2 | | | 7.26 | 0 | n.d. | 1399.565 | 933.380 | 1399.563 | 0.002 |
| H5N4S2F1 | | | 9.26 | 0 | n.d. | 1472.086 | 981.726 | 1472.084 | 0.002 |
| H6N4S1F1 | | | 0.21 | 0 | n.d. | 1407.563 | n.d. | 1407.560 | 0.004 |
| H3N5F1 | | | 4.52 | 3.08 | 2239.937 | 1120.474 | n.d. | 1120.473 | 0.002 |
| H4N5F1 | | | 1.96 | 5.02 | n.d. | 1201.501 | 801.337 | 1201.498 | 0.003 |
| H4N5F2 | | | 3.85 | 0 | 2548.049 | 1274.531 | n.d. | 1274.527 | 0.003 |
| H4N5S1F1 | | | 4.11 | 0 | n.d. | 1346.540 | n.d. | 1346.542 | 0.002 |
| H4N5S2F1 | | | 3.53 | 0 | n.d. | 1492.598 | 995.400 | 1492.594 | 0.004 |
| H5N5F1 | | | 1.40 | 5.40 | n.d. | 1282.529 | 855.354 | 1282.524 | 0.004 |
| H5N5F2 | | | 1.32 | 0 | n.d. | 1355.558 | n.d. | 1355.554 | 0.004 |
| H5N5F3 | | | 0.56 | 0.24 | n.d. | 1428.587 | 952.727 | 1428.583 | 0.004 |
| H5N5S1F1 | | | 3.49 | 0.25 | n.d. | 1428.076 | 952.387 | 1428.072 | 0.004 |
| H6N5F1 | | | 0.78 | 17.15 | n.d. | 1363.555 | 909.372 | 1363.552 | 0.003 |
| H6N5F2 | | | 1.51 | 0 | n.d. | 1436.585 | 958.059 | 1436.580 | 0.005 |
| H6N5S1F1 | | | 7.08 | 1.36 | n.d. | 1509.102 | 1006.405 | 1509.100 | 0.003 |
| H6N5S1F2 | | | 3.21 | 0 | n.d. | 1582.132 | 1055.091 | 1582.129 | 0.004 |
| H3N6F1 | | | 1.19 | 0.70 | n.d. | 1222.014 | 814.676 | 1222.011 | 0.003 |
| H3N6F2 | | | 2.04 | 0 | n.d. | 1295.044 | 863.359 | 1295.042 | 0.002 |
| H3N6S1F1 | | | 1.39 | 0 | n.d. | 1367.564 | 911.706 | 1367.556 | 0.007 |
| H4N6F1 | | | 0.81 | 2.37 | n.d. | 1303.042 | n.d. | 1303.038 | 0.004 |
| H4N6F2 | | | 0.59 | 0 | n.d. | 1376.071 | n.d. | 1376.069 | 0.002 |
| H5N6F1 | | | 0.58 | 2.88 | n.d. | 1383.567 | n.d. | 1383.561 | 0.006 |
| H5N6F2 | | | 1.02 | 0 | n.d. | 1457.098 | 971.733 | 1457.094 | 0.004 |
| H6N6F1 | | | 0 | 5.67 | n.d. | 1464.581 | 977.064 | 1464.586 | 0.005 |
| H6N6S2F1 | | | 0.83 | 0 | n.d. | 1756.697 | 1171.131 | 1756.686 | 0.012 |
| H6N6S3F1 | | | 0.43 | 0 | n.d. | 1901.742 | 1268.162 | 1901.736 | 0.006 |
| H7N6F1 | | | 0.17 | 15.83 | n.d. | 1545.619 | 1030.405 | 1545.614 | 0.005 |
| H7N6S1F1 | | | 0.50 | 0.91 | n.d. | 1691.669 | 1128.116 | 1691.666 | 0.003 |
| H6N7 | | | 0 | 0.60 | n.d. | 1493.595 | 996.065 | 1493.605 | 0.011 |
| H6N7F1 | | | 0 | 0.21 | n.d. | 1566.132 | 1044.757 | 1566.131 | 0.001 |
| H7N7F1 | | | 0 | 1.92 | n.d. | 1647.158 | 1098.776 | 1647.152 | 0.006 |
| H8N7F1 | | | 0 | 8.63 | n.d. | 1728.688 | 1152.793 | 1728.684 | 0.003 |
| H8N7S1 | | | 6.13 | 0 | n.d. | 1800.201 | 1200.810 | 1800.199 | 0.002 |
| H8N8F1 | | | 0 | 0.24 | n.d. | 1829.724 | 1220.152 | 1829.723 | 0.002 |
| H9N8F1 | | | 0 | 2.39 | n.d. | 1911.753 | 1274.505 | 1911.751 | 0.002 |
| H10N9F1 | | | 0 | 0.31 | n.d. | 2093.323 | 1395.881 | 2093.319 | 0.005 |
